# Exome sequencing of 457 autism families recruited online provides evidence for novel ASD genes

**DOI:** 10.1101/516625

**Authors:** Pamela Feliciano, Xueya Zhou, Irina Astrovskaya, Tychele N. Turner, Tianyun Wang, Leo Brueggeman, Rebecca Barnard, Alexander Hsieh, LeeAnne Green Snyder, Donna M. Muzny, Aniko Sabo, The SPARK Consortium, Richard A. Gibbs, Evan E. Eichler, Brian J. O’Roak, Jacob J. Michaelson, Natalia Volfovsky, Yufeng Shen, Wendy K. Chung, Leonard Abbeduto, John Acampado, Andrea J. Ace, Charles Albright, Michael Alessandri, David G. Amaral, Alpha Amatya, Robert D. Annett, Ivette Arriaga, Irina Astrovskaya, Ethan Bahl, Adithya Balasubramanian, Nicole Bardett, Rebecca A. Barnard, Asif Bashar, Arthur Beaudet, Landon Beeson, Raphael A. Bernier, Elizabeth Berry-Kravis, Stephanie Booker, Stephanie J. Brewster, Elizabeth Brooks, Leo Brueggeman, Martin E. Butler, Eric M. Butter, Kristen Callahan, Alexies Camba, Sarah Carpenter, Nicholas Carriero, Lindsey A. Cartner, Ahmad S. Chatha, Wubin Chin, Wendy K. Chung, Renee D. Clark, Cheryl Cohen, Eric Courchesne, Joseph F. Cubells, Mary Hannah Currin, Amy M. Daniels, Lindsey DeMarco, Megan Y. Dennis, Gabriel S. Dichter, Yan Ding, Huyen Dinh, Ryan Doan, HarshaVardhan Doddapaneni, Evan E. Eichler, Sara Eldred, Christine Eng, Craig A. Erickson, Amy Esler, Ali Fatemi, Pamela Feliciano, Gregory Fischer, Ian Fisk, Eric J. Fombonne, Emily A. Fox, Sunday Francis, Sandra L. Friedman, Swami Ganesan, Michael Garrett, Vahid Gazestani, Madeleine R. Geisheker, Jennifer A. Gerdts, Daniel H. Geschwind, Richard A. Gibbs, Robin P. Goin-Kochel, Anthony J.Griswold, Luke P. Grosvenor, Angela J. Gruber, Amanda C. Gulsrud, Jaclyn Gunderson, Anibal Gutierrez, Melissa N. Hale, Monica Haley, Jacob B. Hall, Kira E. Hamer, Bing Han, Nathan Hanna, Christina Harkins, Nina Harris, Brenda Hauf, Caitlin Hayes, Susan L. Hepburn, Lynette M. Herbert, Michelle Heyman, Brittani A. Hilscher, Susannah Horner, Alexander Hsieh, Jianhong Hu, Lark Y. Huang-Storms, Hanna Hutter, Dalia Istephanous, Suma Jacob, William Jensen, Mark Jones, Michelle Jordy, A. Pablo Juarez, Stephen Kanne, Hannah E. Kaplan, Matt Kent, Alex Kitaygorodsky, Tanner Koomar, Viktoriya Korchina, Anthony D. Krentz, Hoa Lam Schneider, Elena Lamarche, Rebecca J. Landa, Alex E. Lash, J. Kiely Law, Noah Lawson, Kevin Layman, Holly Lechniak, Sandra Lee, Soo J. Lee, Daniel Lee Coury, Christa Lese Martin, Hai Li, Deana Li, Natasha Lillie, Xiuping Liu, Catherine Lord, Malcolm D. Mallardi, Patricia Manning, Julie Manoharan, Richard Marini, Gabriela Marzano, Andrew Mason, Emily T. Matthews, James T. McCracken, Alexander P. McKenzie, Jacob J. Michaelson, Zeineen Momin, Michael J. Morrier, Shwetha Murali, Donna Muzny, Vincent J. Myers, Jason Neely, Caitlin Nessner, Amy Nicholson, Kaela O’Brien, Eirene O’Connor, Brian J. O’Roak, Cesar Ochoa-Lubinoff, Jessica Orobio, Opal Y. Ousley, Lillian D. Pacheco, Juhi Pandey, Anna Marie Paolicelli, Katherine G. Pawlowski, Karen L. Pierce, Joseph Piven, Samantha Plate, Marc Popp, Tiziano Pramparo, Lisa M. Prock, Hongjian Qi, Shanping Qiu, Angela L. Rachubinski, Kshitij Rajbhandari, Rishiraj Rana, Rick Remington, Catherine E. Rice, Chris Rigby, Beverly E. Robertson, Katherine Roeder, Cordelia R. Rosenberg, Nicole Russo-Ponsaran, Elizabeth Ruzzo, Aniko Sabo, Mustafa Sahin, Andrei Salomatov, Sophia Sandhu, Susan Santangelo, Dustin E. Sarver, Jessica Scherr, Robert T. Schultz, Kathryn A. Schweers, Swapnil Shah, Tamim Shaikh, Yufeng Shen, Amanda D. Shocklee, Andrea R. Simon, Laura Simon, Vini Singh, Steve Skinner, Christopher J. Smith, Kaitlin Smith, LeeAnne G. Snyder, Latha V. Soorya, Aubrie Soucy, Alexandra N. Stephens, Colleen M. Stock, James S. Sutcliffe, James S. Sutcliffe, Amy Swanson, Maira Tafolla, Nicole Takahashi, Carrie Thomas, Taylor Thomas, Samantha Thompson, Jennifer Tjernagel, Tychele N. Turner, Bonnie Van Metre, Jeremy Veenstra-Vanderweele, Brianna M. Vernoia, Natalia Volfovsky, Jermel Wallace, Corrie H. Walston, Jiayao Wang, Tianyun Wang, Zachary Warren, Lucy Wasserburg, Loran Casey White, Sabrina White, Ericka L. Wodka, Simon Xu, Wha S. Yang, Meredith Yinger, Timothy Yu, Lan Zang, Hana Zaydens, Haicang Zhang, Haoquan Zhao, Xueya Zhou

## Abstract

Autism spectrum disorder (ASD) is a genetically heterogeneous condition, caused by a combination of rare *de novo* and inherited variants as well as common variants in at least several hundred genes. However, significantly larger sample sizes are needed to identify the complete set of genetic risk factors. We conducted a pilot study for SPARK (SPARKForAutism.org) of 457 families with ASD, all consented online. Whole exome sequencing (WES) and genotyping data were generated for each family using DNA from saliva. We identified variants in genes and loci that are clinically recognized causes or significant contributors to ASD in 10.4% of families without previous genetic findings. Additionally, we identified variants that are possibly associated with autism in an additional 3.4% of families. A meta-analysis using the TADA framework at a false discovery rate (FDR) of 0.2 provides statistical support for 34 ASD risk genes with at least one damaging variant identified in SPARK. Nine of these genes (*BRSK2, DPP6, EGR3, FEZF2, ITSN1, KDM1B, NR4A2, PAX5* and *RALGAPB*) are newly emerging genes in autism, of which *BRSK2* has the strongest statistical support as a risk gene for autism (TADA q-value = 0.0015). Future studies leveraging the thousands of individuals with ASD that have enrolled in SPARK are likely to further clarify the genetic risk factors associated with ASD as well as allow accelerate autism research that incorporates genetic etiology.

## Introduction

Autism spectrum disorder (ASD) is an extremely variable condition characterized by deficits in social interactions and restrictive, repetitive behaviors. Currently, there are no FDA approved medications that address these core symptoms, despite the life-long morbidity and increased mortality in adults with autism^1^.

Despite the significant clinical heterogeneity of this condition, many studies have shown that ASD is highly heritable, with genetic risk factors thought to explain the majority of the risk for ASD^2^. Over the past decade, genomic studies focused on recurrent, *de novo*, likely gene disrupting (dnLGD) variants (stopgain, frameshift, and essential splice site) have identified ~100 high-confidence autism risk genes or loci^3–4^. Previous studies have identified molecular diagnoses in 6-37% of individuals with ASD, with higher yields in individuals with additional comorbidities that include intellectual disabilities, seizures, and other medical features^5^.

Here we describe the results of a pilot study that genetically characterized 457 families with one or more members affected with ASD enrolled online in SPARK^6^. SPARK’s mission is to create the largest recontactable research cohort of at least 50,000 families affected with ASD in the United States for longitudinal phenotypic and genomic characterization who are available to participate in research studies. Using exome sequencing and genome-wide single nucleotide polymorphism (SNP) genotyping arrays, we identified variants that are the likely primary genetic cause of autism in 14% of families. We also demonstrated that the genetic architecture in this self-reported cohort is similar to published, clinically-confirmed autism cohorts^3,7,4^. Combining the SPARK data with prior studies, our analyses provide strong evidence that *BRSK2* is a new high-confidence ASD risk gene (FDR q-value = 0.0015) and provide additional evidence for eight newly emerging risk genes (*DPP6, EGR3, FEZF2, ITSN1, KDM1B, NR4A2, PAX5* and *RALGAPB*) in ASD.

## Results

### Variant discovery

We report the exome sequencing and genotyping results of 1379 individuals in 457 families with at least one offspring affected with ASD, including 418 simplex and 39 multiplex families (Supplementary Figure 1). Over 80% of participants are predicted to have European ancestry based on principal component analysis of common SNP genotypes (Supplementary Figure 2). The male to female ratio is 4.4:1 among 418 cases in simplex families, and 2.9:1 among 47 offspring in multiplex families. Of the 465 offspring affected with ASD, 25.6% also reported intellectual disability (Table 1). We identified 647 rare (allele frequency AF<0.001 in ExAC v0.3) *de novo* single nucleotide variants (SNVs) and indels (Supplementary Table 1) in coding regions and splice sites (1.4/offspring), including 85 likely gene disrupting (LGD) variants and 390 missense variants. Similar to the *de novo* variants identified from 4,773 clinically ascertained ASD trios from previous studies^3–8^, the frequency of dnLGD variants in the 465 affected offspring (0.18/offspring) is 1.76-fold higher than the baseline expectation calculated by a previously published mutation rate model^9^ (p = 1.2 × 10^−6^ by one-sided exact Poisson test) (Methods; Supplementary Table 2A).

To identify *de novo* missense variants that are likely damaging, we applied two deleterious missense (D-mis) prediction algorithms on published ASD and SPARK *de novo* variants. Among the 390 *de novo* missense variants in affected offspring, 43.6% are predicted to be deleterious using CADD ≥ 25^10^ and show 1.28-fold enrichment compared to baseline expectation in the general population. Using a stricter D-mis prediction algorithm with MPC≥2^11^, eight percent of *de novo* missense variants are predicted as deleterious and are enriched 1.88-fold in affected offspring which is comparable to the enrichment of dnLGD variants. The overall burden of *de novo* D-mis variants is similar to published studies (Supplementary Table 2B).

**Table 1:**
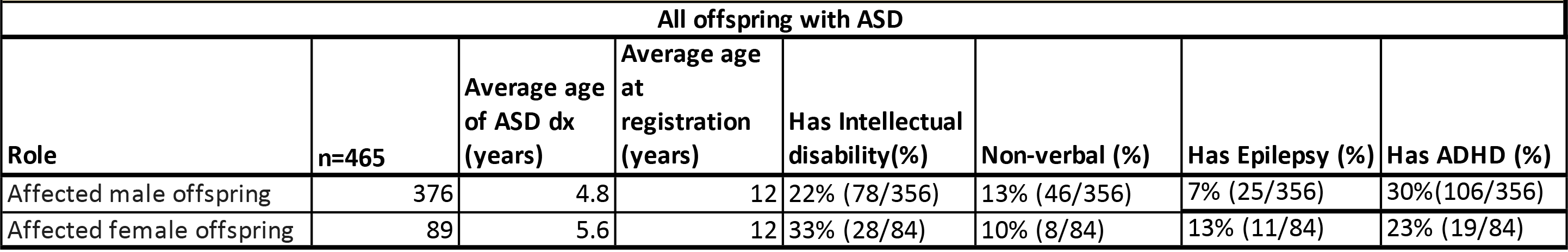
Phenotypic description of the 457 families with at least 1 offspring affected with ASD in the SPARK pilot study. 1379 individuals in 39 multiplex and 418 simplex families were genomically characterized, including 472 individuals (465 offspring and 7 parents affected with ASD) were sequenced. Phenotypic variables are not available for each person, so they are reported for whom they are available.

| All offspring with ASD |  |  |  |  |  |  |  |
| --- | --- | --- | --- | --- | --- | --- | --- |
| Role | n=465 | Average age of ASD dx (years) | Average age at registration (years) | Has Intellectual disability(%) | Non-verbal (%) | Has Epilepsy (%) | Has ADHD (%) |
| Affected male offspring | 376 | 4.8 | 12 | 22% (78/356) | 13% (46/356) | 7% (25/356) | 30%(106/356) |
| Affected female offspring | 89 | 5.6 | 12 | 33% (28/84) | 10% (8/84) | 13% (11/84) | 23% (19/84) |

Variants in constrained genes (pLI ≥ 0.5)^12^ explain most of the burden of dnLGD variants and *de novo* D-mis variants (defined by a MPC score ≥ 2) in the affected offspring in our study (Supplementary Table 2B). Consistent with previous findings supporting the female protective model^13^, we observed a non-significant trend toward a higher frequency of dnLGD variants in constrained genes in female cases compared to males (0.135/female vs. 0.096/male), as well as higher frequency of *de novo* D-mis variants in female cases (CADD=25: 0.416/female vs 0.354/male, MPC ≥ 2: 0.09/female vs 0.066/male).

We also investigated deleterious inherited SNV/indel variants and found a modest excess of transmitted, rare LGD (AF < 0.001 in ExAC v0.3) variants observed only once among parents in our cohort (singletons) in constrained genes with pLI ≥ 0.5 (464 transmitted vs. 402 non-transmitted; RR = 1.15, P = 0.038 by binomial test). Over-transmission of rare singleton LGD variants was not observed in genes that are not constrained (RR=1.03, P=0.31 by binomial test). The excess of transmitted singleton LGD variants in constrained genes increased after removing variants observed in the ExAC database (303 transmitted vs. 242 un-transmitted; RR = 1.25, P = 0.010 by binomial test). These results provide further evidence that rare, inherited LGD variants in constrained genes confer increased risk for ASD^14,15^. We then searched for known haploinsufficient ASD or neurodevelopmental disorder (NDD) genes (SFARI gene score 1 or 2 or listed in DDG2P^16^) that are disrupted by the rare singleton LGD variants and are transmitted. We found 13 such variants (2 of them on the X chromosome), as compared to 10 variants that are not transmitted (including one on the X chromosome) (Supplementary Table 3). Manual review of these variants revealed that most of them are not likely pathogenic because they either affect only a subset of transcripts that are not expressed in the majority of tissues^17^, are located close to the 5’ end of the transcript (last 5% of the coding sequence) or are indels that overlap but do not change the sequence of essential splice sites. The results suggest that the rare LGD variants in known ASD/NDD genes have only limited contribution to the overall transmission disequilibrium in this class of variants.

By integrating exome sequence read depth and SNP microarray signal intensity data, we identified 273 rare CNVs (occurring with an allele frequency of ≤1% of the 1379 individuals in the analysis) in 206 affected offspring. Of these, 253 CNVs were inherited (0.544/affected offspring) and were on average 194 kb. These inherited CNVs contained an average of 4 genes, which reduces to an average of 0.4 genes that are highly intolerant to variation (pLI ≥ 0.9) (Supplementary Table 4). Similar to the frequency observed in previous studies^8,18^ (0.052/proband), we identified 20 *de novo* CNVs (0.043 /affected offspring) (Supplementary Table 5). On average, *de novo* CNVs were larger (1.6 Mb) and contained more total and constrained genes (28 genes total, 3 genes with a pLI ≥ 0.9).

Despite the fact we were underpowered to detect statistically significant burden differences between sexes, we observed a trend toward a 1.8-fold higher burden of dnCNVs in ASD females (0.067/female vs. 0.037/male, respectively). In contrast, the frequency of rare, inherited CNVs in ASD females and males were similar (0.551/female vs. 0.543/male, respectively). Similar to Sanders et al. 2015^8^, dnCNVs in female cases also affect more genes than dnCNVs in males (0.669 vs. 0.039 genes in dnCNVs per female proband vs per male proband, respectively; p = 0.013, Kruskal-Wallis test).

Of the CNVs detected, there were six mapping within the chromosome 16p11.2 region (3 *de novo* and 3 inherited in five families). Four of the six 16p11.2 CNVs occurred at the most common breakpoints (BP4-BP5), occurring in 0.9% of affected offspring, consistent with the expected autism prevalence^19^. Together, the results suggest that the saliva-derived DNA collected in SPARK should provide comparable CNV data to previous studies using DNA derived from whole blood. We also used read-depth and SNP genotypes to identify several chromosomal aneuploidies (Supplementary Figure 1B, including one case of trisomy 21 (47, XY +21), one case of Klinefelter syndrome (47, XXY), one case of Turner syndrome (45, X), and one case of uniparental iso-disomy of chromosome 6 (UPiD6).

Given their emerging role in genetic risk for ASD and other NDDs, we also assessed postzygotic mosaic mutations^20–21^ in the SPARK cohort. In parallel, we utilized a previously established method^22^ and a novel approach to identify likely mosaic SNVs (Methods, Supplementary Figures 3-8). We identified 65 likely mosaic mutations (0.142/offspring) (Supplementary Table 6). The majority of these mutations were unique to the mosaic call set; however, 18 were also identified in the main *de novo* SNV call set with an average alternative allele fraction of 25.4% (Supplementary Table 6), suggesting that these mutations are likely to have occurred post-fertilization. These results indicate that ~10% (65/652) of the total *de novo* SNVs in the SPARK pilot are of postzygotic origin. Comparing these data to a similar mosaic set from the SSC^22^, we found similar mosaic mutation characteristics, despite the fact that different DNA sources, capture reagents, and sequencing instruments were used (Supplementary Figure 7). Due to the limited number of mosaic calls, we did not attempt to evaluate mosaic mutation burden. However, we observed that a number of potentially mosaic mutations were in known or candidate risk genes ASD/NDD or genes that are highly constrained (Supplementary Table 6). For example, we identified a mosaic LGD variant in *MACF1*, which is highly constrained (pLI =1), plays essential roles in neurodevelopment, functions through the previously implicated Wnt signaling pathway^23^, and has been recently suggested as a candidate gene based on a *de novo* LGD variant in a Japanese ASD cohort^24^. In *CREBBP*, which reached genome-wide significance in a recent NDD meta-analysis^16^, we identified a mosaic missense variant, in addition to two other germline *de novo* missense variants in SPARK, adding to the evidence that it is an ASD/NDD risk factor.

### Genes with a higher mutational burden

We assessed genes with recurrent *de novo* LGD variants in the SPARK cohort and identified four genes with more than one *de novo* LGD variant (*CHD8*, *FOXP1*, *SHANK3*, *BRSK2*). *BRSK2* is the only gene with multiple *de novo* LGD variants in SPARK and not previously implicated in autism and/or NDDs (p=2.3 ×10^−6^ by one-sided exact Poisson test).

To increase the statistical power to identify new autism genes, we performed a meta-analysis of *de novo* variants in 4,773 published ASD trios^3,4,7,8^ and 470 SPARK trios using TADA^25^ (Methods). In this analysis, we included *de novo* LGD variants and *de novo* D-mis variants, which we defined as those that have a CADD score ≥ 25 ^10^. The TADA analysis presumes a model of genetic architecture compatible with the observed burden and recurrence of *de novo* damaging variants and assigns an FDR q-value for each gene based on the number of damaging variants and baseline mutation rates. We identified 34 genes with at least one damaging variant identified in SPARK, which also met our FDR threshold (< 0.2) (Figure 1). In our TADA results, we only present genes with damaging variants in the SPARK data (Supplementary Table 7). Our simulations (Supplementary Table 8) show that by limiting results to variants observed in the SPARK cohort, the FDR threshold will be conservative among those reported genes (Supplementary Note). Restricting the TADA analysis to only the published *de novo* variants from 4,773 trios, 16 known ASD/NDD genes were significant at FDR<0.2 while the other 18 genes were non-significant (FDR > 0.4). For the latter 18 genes, previous studies identified a small number of *de novo* damaging variants. The additional damaging variants identified in the 457 families from the SPARK cohort were sufficient to significantly strengthen the statistical evidence in the combined cohort. We also performed TADA analyses including inherited variants and CNVs from the SPARK families, but we found that these variant classes do not contribute to the statistical evidence for the variants identified above.

**Figure 1:**
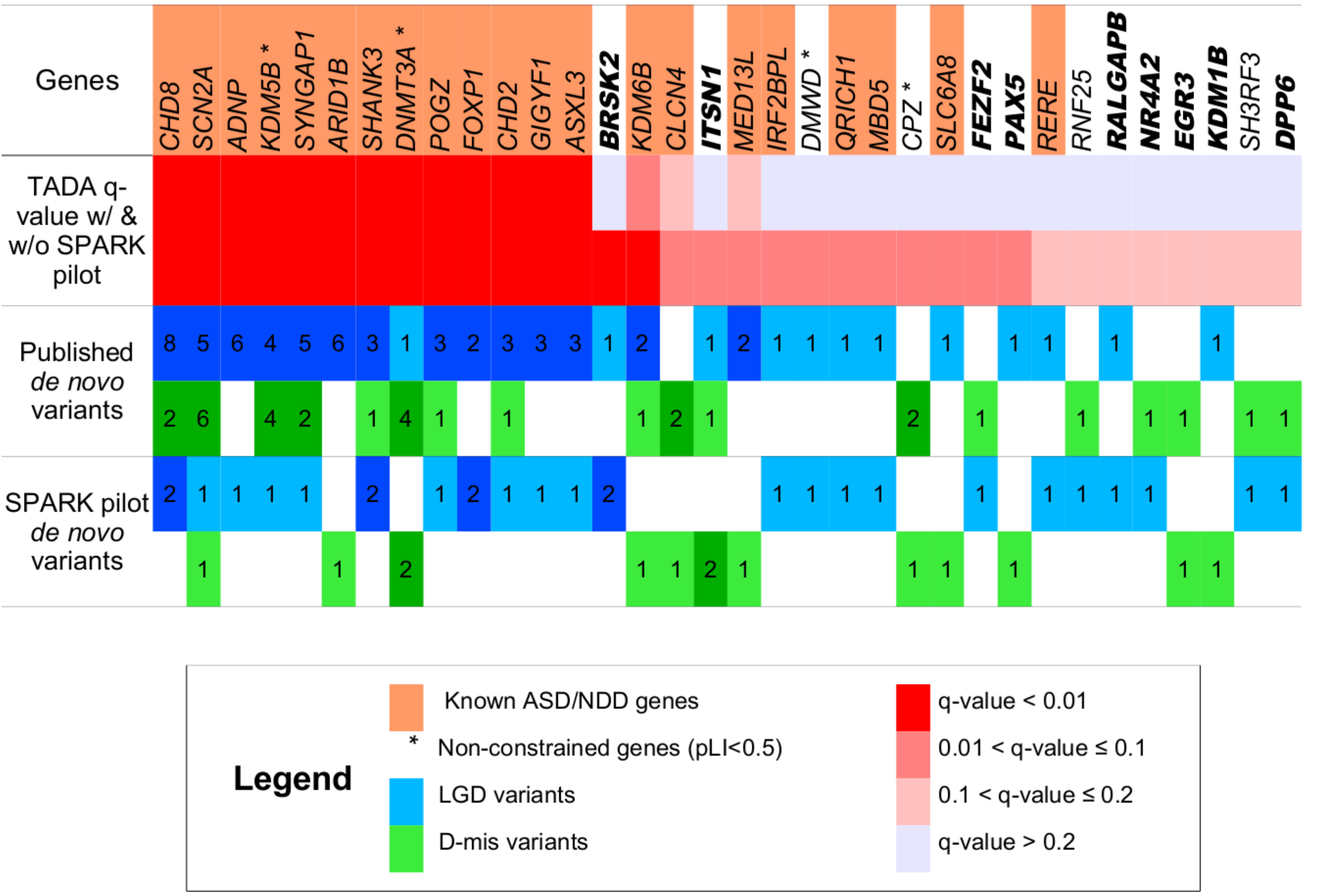
Meta-analysis using the TADA framework provides statistical support for 34 genes at a false discovery rate (FDR) of 0.2. Known ASD genes are defined as those with SFARIgene score ≤ 2 or implicated in a previous TADA meta-analysis (FDR<0.1)8; known NDD genes are those listed in the DDG2P database16. D-mis variants are defined by CADD score ≥ 25. A total of 34 genes with at least one de novo damaging variant observed in SPARK pilot trios achieve an FDR<0.2 after meta-analysis with published trios (total n=5,230). Fourteen genes have not been previously classified as known ASD or NDD genes. Nine of these genes are supported by additional evidence (Supplementary Table 10) and are considered newly emerging ASD risk genes.

Of the newly identified genes, *BRSK2* has the strongest statistical evidence as a new autism gene, with a q-value = 0.0015. All four individuals in SPARK, ASC and the SSC with *de novo* functional variants in *BRSK2* are males with cognitive impairment and severe speech delay (Table 2). Among these 18 genes, 16 are constrained genes (pLI ≥ 0.5). Since the association signal for the two non-constrained genes was driven by LGD variants, we excluded these from further analysis as likely false-positives. *MBD5* and *IRF2BPL* achieved a FDR value of < 0.1 in a previous meta-analysis^8^, which also included evidence from *de novo* CNVs and deleterious variants of unknown inheritance from a case-control sample in that analysis. Four genes (*QRICH1*, *MBD5*, *SLC6A8*, and *RERE*) are known NDD risk genes in the latest DDG2P database^16^.

We then searched for additional supporting evidence for a role of these genes in autism and NDDs, including other deleterious variants in previous studies and case reports not included in the meta-analysis, membership in gene sets previously associated with ASD^3,4,7,8^, and published functional studies (Supplementary Table 9). Previous studies have reported additional individuals with NDDs and/or autism with *de novo* damaging variants in four genes (*PAX5*^4,26^, *NR4A2*^27,28^*, RALGAPB*^7,29,30^, and *DPP6*^5,31,32^).

In addition to recurrent deleterious variants in these newly emerging ASD risk genes, we also found evidence that they function in biological pathways previously linked to autism. For example, mRNA translation of *BRSK2, ITSN1,* and *RALGAPB* in neurons is predicted to be regulated by FMR1 protein^33^. In addition, *ITSN1* and *DPP6* are part of the post-synaptic density components in human neocortex^34^. *PAX5* and *FEZF2* are involved in transcription regulation during central nervous system development^4,24,35^. *KDM1B* is a known chromatin modifier, and *EGR3* has been implicated in neurodevelopment^36,37^. Combining statistical and functional evidence, our analysis provides additional support for nine genes (*BRSK2, DPP6, EGR3, FEZF2, ITSN1, KDM1B, NR4A2, PAX5* and *RALGAPB*) as newly emerging genes in ASD and NDDs.

We also searched rare singleton inherited LGD variants in these nine genes in SPARK and published Simons Simplex Collection (SSC) data, and identified five additional cases (3 in SSC, 2 in SPARK) carrying inherited LGD variants of *ITSN1* that likely cause loss of gene function. Interestingly, of the six ASD cases with LGD variants in *ITSN1*, five do not have intellectual disability (Table 2). The less severe phenotype and inheritance from unaffected parents are consistent with the modest effect size. Furthermore, in ASC case-control samples^7^, LGD variants in *ITSN1* were also identified in the controls, although they were still over-represented in cases (2 in 1601 cases vs 3 in 5397 controls and comparable to the cumulative AF of 2.5e-4 in gnomAD v2.1). We did not find any rare, inherited variants causing loss of function in other newly emerging genes from affected offspring.

### Functional network analysis and gene expression patterns in newly emerging genes

To relate the newly emerging ASD risk genes from our genetic analysis to previous knowledge of integrated gene networks in ASD, we scored the newly emerging genes using forecASD, a new ensemble classifier that integrates spatiotemporal gene expression, heterogeneous network data, and previous gene-level predictors of autism association^38^. Using forecASD, we derived a single score that ranks the evidence for each gene to be involved in autism risk. Using this approach, we found that the nine newly emerging ASD risk genes are all in the top decile of forecASD scores (the top decile being a recommended cutoff used to define probable ASD risk genes), and correspondingly have significantly elevated forecASD scores over the remainder of the genome (P=8.8×10^−7^, Supplementary Figure 9), supporting these genes as having similar properties overall compared to known ASD risk genes. Furthermore, two predictive features in forecASD that summarize brain expression support and network support are also found to be significantly elevated over the genome background in the new genes (P=0.017 and P=0.004, respectively; see Supplementary Figure 9). Importantly, neither of these metrics uses genetic data directly, so the nine newly emerging genes collectively have support across the three independent and distinct domains of genetic, network, and brain expression evidence.

**Table 2:**
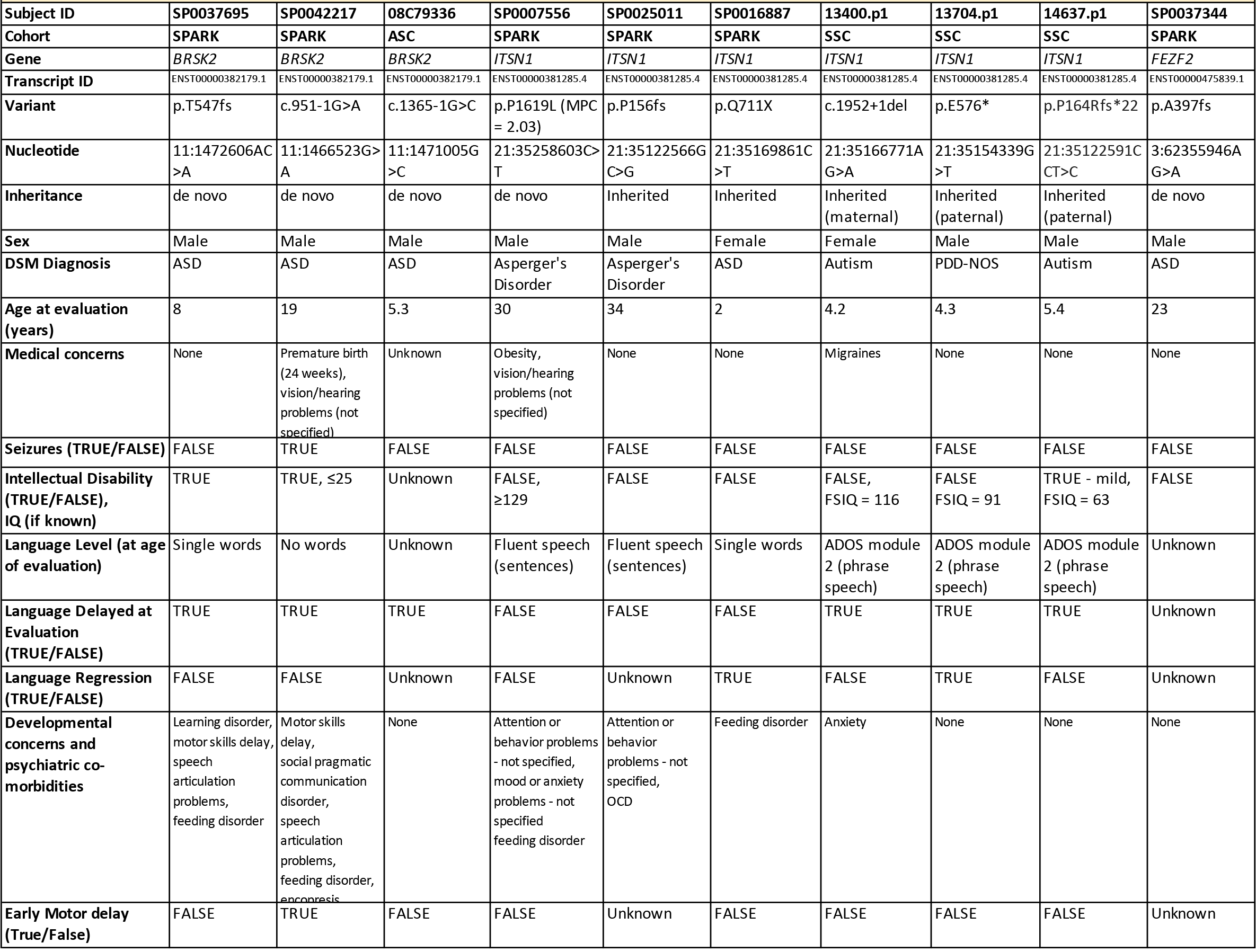
Phenotypic information on individuals with deleterious variants in the nine newly emerging ASD risk genes. MPC^11^ scores are listed for missense mutations. All phenotypic information for SPARK participants was collected online. Because phenotypic data collection procedures are not consistent between different research cohorts, we implemented a systematic, quantitative phenotypic severity rating scale based on consistent variables between cohorts. This rating scale incorporated indicators of intellectual disability, birth defects and presence of seizures and allows for easier comparison of phenotypes between individuals in different cohorts (Methods). Severity is scored on a scale of 0 to 8, with 8 indicating most severe and 0 indicating unremarkable.

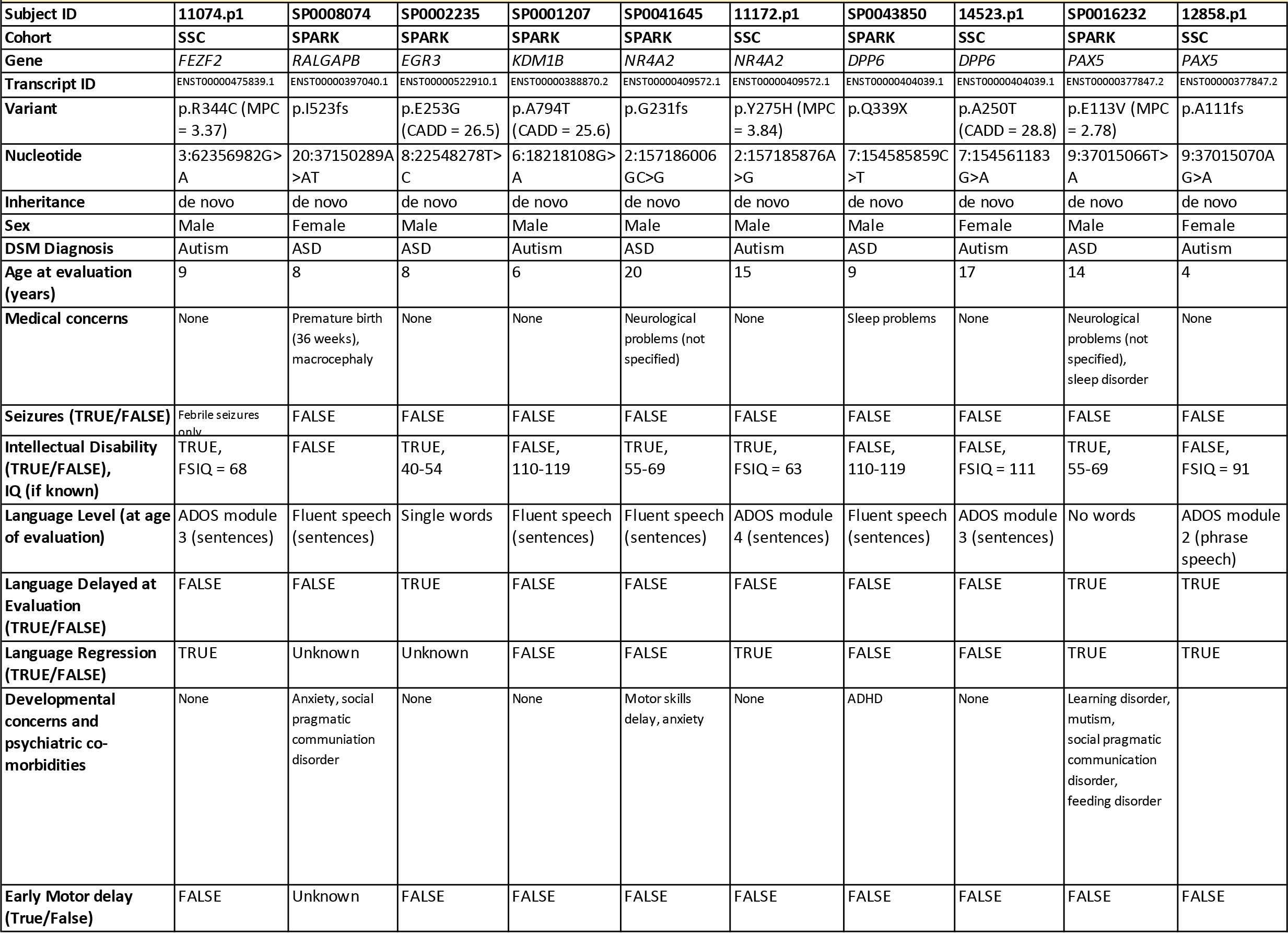

To illustrate the network context of the newly emerging ASD risk genes, we clustered the new genes along with known ASD risk genes (SFARI score 1 or 2) and genes scoring within the top decile of forecASD (Figure 2A). Network analysis yielded 11 tightly connected clusters with distinct biological functions (Supplementary Table 10). Several genes were assigned to clusters that showed enrichment for gene sets consistent with their known functions, including *DPP6*^39^, *KDMB1*^40^, *NR4A2*^41^, and *FEZF2*^42^, consistent with their functional evidence (Supplementary Table 10). In a subsequent analysis, the interactions between known ASD risk genes and the nine newly implicated ASD risk genes were visualized (Figure 2B). This subnetwork was significantly interconnected (P<1 × 10^−16^), with novel genes showing significantly more functional associations with known ASD risk genes than expected by chance (P<0.001).

Using coexpression networks seeded by high-confidence ASD risk genes, a previous study found that cortical projection neurons in layers 5 and 6 of human midfetal prefrontal and primary motor-somatosensory cortex (PFC-MSC) are a key point of convergence for ASD risk genes^43^. Another study also showed that unbiased gene co-expression networks overrepresented with candidate ASD risk genes are more highly expressed in the cortical plate and subplate laminae of the developing human cortex, which will go on to form mature layers II-VI of the cerebral cortex^44^. One of the newly emerging genes we identified, *FEZF2*, is a powerful master regulator gene critical for establishing corticospinal neurons^45^, which connect layer Vb of the cortex to the spinal cord, and is known to be expressed in the putative layer V in the late mid-fetal human cortex^46^.

We first evaluated gene expression of the newly emerging genes with regard to cortical layer specificity in the human developing brain^47^. Among the nine newly emerging genes, seven (*BRSK2, ITSN1, FEZF2, RALGAPB, NR4A2, EGR3* and *DPP6*) have expression data in developing fetal human cortex, and similar to Parikshak et al.^44^, they show a trend of increased expression at post-conceptual week (PCW) 15-16 (Figure 2c) and PCW 21 (Supplementary Figure 10) in the cortical plate and subplate laminae, which will form layers II-VI of the mature cerebral cortex. The mean of t-statistics of these seven genes in the inner cortical plate (CPi) and subplate (SP) are greater than two standard deviations (SD) from the mean of randomly selected genes matched for gene length and GC content (P < 0.01 by simulation).

We further evaluated cell type specificity using recently published single-cell RNA-seq data from fetal and adult mouse and human brains^48^ (Supplementary Figures 11-12), and found the expression specificity of newly implicated genes is highest in pyramidal neurons in the mouse hippocampus CA1 region with an enrichment of 4.3 SD from the bootstrapped mean (p=5.6e-3 by simulations controlling gene length and GC content, Supplementary Figure 11). The specificity is also enriched (2.1 SD above the bootstrapped mean, p=0.03 by simulation) in somatosensory pyramidal neurons using a recently published human single nucleus RNA-seq data^49^. These results are consistent with a previous study showing that autism protein-protein interaction networks related to the 16p11.2 CNV display significantly enriched expression during mid-fetal development as well as early childhood in cerebral cortex^50^. Taken together, we find that the newly emerging genes identified in this study demonstrate differential expression patterns similar to that of known ASD genes, providing further support that the newly emerging genes function in similar biological pathways and mechanisms as known ASD genes.

**Figure 2:**
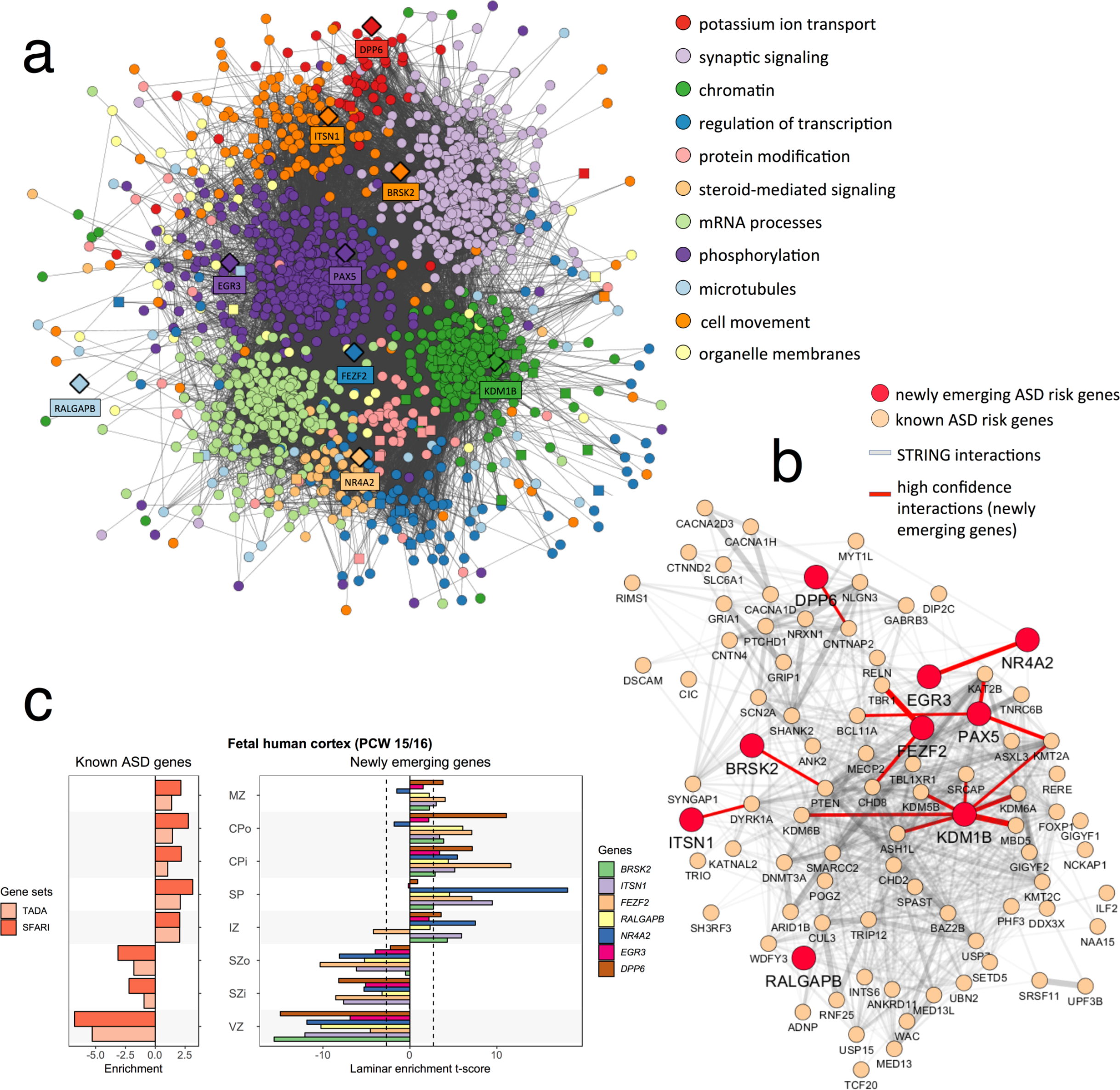
Network analysis & gene expression of newly emerging ASD genes using A) GO term clustering, B) PPI networks and C) gene expression of human fetal cortex at PCW (post-coital weeks) 15-16. Known ASD genes are defined as those with a SFARIgene score ≤2 (84 genes, indicated as SFARI) or implicated in a previous TADA meta-analysis at FDR<0.1 (65 genes, indicated as TADA). The enrichment for each gene was measured by the t-statistics comparing the expression level in each layer against all other layers. The enrichment of a gene set is the mean of t-statistics of its genes. Two newly emerging ASD risk genes (PAX5 and KDM1B) are not shown due to the low expression levels in human developing cortex (RPKM<1 for at least 20% available neocortical samples in BrainSpan). Data were extracted from Supplementary Table of 44. Laminae abbreviations: marginal zone (MZ), outer/inner cortical plate (CPo/CPi), subplate (SP), intermediate zone (IZ), outer/inner subventricular zone (SZo/SZi), ventricular zone (VZ).

### Diagnostic yield in SPARK

For all cases, we defined deleterious autism-associated variants as those meeting likely pathogenic or pathogenic criteria according to ACMG standards^51^ (Methods). We did not search for or discover any incidental findings unrelated to autism in these families. We defined possible autism-associated variants as either: SNVs that are *de novo* missense variants that affect known NDD or ASD genes and have an MPC score^11^ ≥ 2, loss-of-function variants that disrupt possible NDD or ASD genes, or CNVs that delete one or more possible NDD or ASD genes or duplicate known ASD or NDD loci. Families in the pilot study were selected without regard to genetic diagnosis. 13 of the 457 families self-reported a genetic diagnosis, and all were confirmed by our analyses and serve as positive controls to validate our genomic analyses (Supplementary Table 11). For the remaining 444 families, we identified 50 (10.8%) deleterious genetic variants (8 *de novo* CNVs, 14 inherited CNVs, 23 *de novo* SNVs or indels, 3 inherited LGD variants and 2 chromosomal aneuploidies) in known autism-associated genes or loci in 49 affected individuals (Supplementary Table 11). We also identified 19 more likely deleterious genetic variants (one *de novo* CNV, one inherited CNV, 14 *de novo* SNVs and three inherited SNVs) in possibly autism-associated genes or loci in an additional 14 individuals (3.1%). When DNA was available, autism-associated genetic findings were confirmed by Sanger sequencing or chromosome microarray, and genetic results were returned to the families (n=28).

We identified 8 families in which one of the affected children had >1 pathogenic or possibly autism-associated variants of different types (Supplementary Figure 13). Six of the eight affected children (75%) also have intellectual disability, which may provide support for the oligogenic model proposed to explain some of the phenotypic heterogeneity of NDD^52^. In 39 multiplex families, pathogenic variants were identified in 6 affected offspring in 5 families (Supplementary Figure 14). The genetic variants contributing to autism in multiplex families are likely complex^4^, with ≥ 1 contributing genetic risk factor of varying penetrance, especially those without additional medical conditions or congenital anomalies.

## Discussion

Overall, the genomic characterization of 457 autism families (418 simplex and 39 multiplex) in SPARK implicates nine emerging genes in ASD that converge on similar biological networks as known ASD risk genes. We identified a returnable genetic result related to autism in 10.8% of affected offspring and have begun returning individual genetic results to the families after confirming results in a clinical laboratory. Not surprisingly, our diagnostic yield was highest in affected individuals who also report presence of seizures (27%). The yield in individuals who also report intellectual disability was also higher (20%) than the overall cohort.

In our analysis, our diagnostic yield in affected offspring in multiplex families (15.2%) was slightly higher than affected offspring in simplex families (10.1%). Interestingly, the genetic findings in multiplex families rarely explained autism in all affected family members. For example, in a family with an affected father and 3 affected children, the most severely affected child harbored a *de novo* LGD in *ADNP*. No other family member carried this variant or any other identifiable contributing variant. In another pedigree, an affected male child with an affected male father inherited a 15q11.2 BP1-BP2 deletion from a mother who does not report an ASD diagnosis, but we found no contributing variant in the affected father. We also identified eight families in which there was >1 contributing variant, even in families in which we were unable to identify contributing variants in all affected offspring. In one family with two affected children, the female child inherited a 1q21.1 CNV from an unaffected mother and also harbored a *de novo* LGD in *RALGAPB*. However, the affected male child did not harbor either of these variants. Future studies with larger sample sizes will allow for a more robust comparison of the genetic architecture of ASD in simplex vs. multiplex families.

Over time, we expect the diagnostic yield in SPARK to increase as more individuals with ASD are studied and as additional genetic risk factors are identified. For example, we identified LGD variants in *CHD3, MEIS2,* and *AKAP10* and deletions of *NFIB, DLL1*, and *HNRNPD* genes. Although these genes did not reach statistical significance in our TADA meta-analysis, their role in ASD is supported by recurrent mutations in the literature, and they likely represent other newly emerging ASD genes (Supplementary Table 9). We interpreted those variants as possible contributors to autism in those individuals. The genetic findings in those cases will be confirmed and returned in the future if and when these genes are established as ASD associated genes.

Using a systems biology approach, we demonstrated that the newly emerging ASD genes identified in this analysis are well-supported beyond genetic association and are predicted to be ASD risk genes based on a variety of functional properties, including patterns of spatiotemporal gene expression in the brain and protein network connectivity. All nine newly emerging genes scored in the top decile of forecASD, an integrator of functional evidence for ASD risk genes (Supplementary Figure 9). The genes localized to network clusters representing processes critical for neurodevelopment (Figure 2), including chromatin modification (*KDM1B*), establishing specific neuronal cell fates^45^ (*FEZF2*), neuronal polarity^53^ (*BRSK2*) and neuronal migration of pyramidal neurons^54^ (*ITSN1*). The newly emerging genes also showed significant over-connectivity to known ASD risk genes (P<1 × 10^−16^, Figure 2B). Together, the TADA genetic association analysis coupled with the supporting functional and network-level data triangulate these genes as being robust and biologically plausible contributors to ASD risk.

Despite the limited sample size in this pilot study, we were able to identify nine newly emerging ASD genes. Power analysis using a simulation-based approach confirmed that the observed yield is expected given the presumed genetic architecture in the TADA analysis (Supplementary Table 12). We expect to identify ~70-75% of all ASD risk genes in the future that meet a similar FDR threshold (0.1- 0.2) when we reach SPARK’s goal of sequencing 50,000 complete trios (Supplementary Table 12). Other analyses of large cohorts in ASD are underway, including a recent analysis of ~12,000 individuals with ASD^55^. This study, which used a mixture of family-based and case-control data, found statistical support for 99 autism risk genes, increasing the number of autism risk genes from 65^8^. Future meta-analyses of both SPARK data and other autism cohort data are planned to maximize autism risk gene discovery.

For most genes identified with *de novo* damaging variants, inherited LGD variants in affected individuals were not found (Kosmicki et al. 2017^15^ and this study), suggesting our current knowledge about ASD risk genes is biased toward those with high penetrance. Future studies with larger sample sizes will be needed to identify and validate additional risk genes of lower penetrance that confer inherited autism risk.

Altogether, these data suggest that the methods used to ascertain individuals with ASD, saliva collection, and genomic data are of high quality, and future analysis of the tens of thousands of families enrolling in SPARK will significantly contribute to our understanding of the genetic basis of ASD. By returning genetic results to participants, we expect to increase engagement and increase the number of recontactable participants for genetically targeted clinical research and trials.

### List of all Display Items (in order of appearance)

**Supplementary Figure 1:** Sample quality controls. **a)** Relatedness was verified based on the scatterplot of the estimated kinship coefficient and number of SNPs with zero shared alleles (IBS0). Parent-offspring, sibling pairs, and unrelated pairs can be distinguished as separate clusters on the scatterplot. One outlier parent-offspring pair (SP0002452 and mother) showed higher than expected IBS0 and was caused by parental chr6 iso-UPD. **b)** Sample sex was verified based on the ratio of heterozygous to homozygous genotypes on the X-chromosome, using normalized sequencing depth of X and Y chromosomes. Individuals with chromosomal abnormalities are highlighted.

**Supplementary Figure 2:** Principal component (PC) analysis of sample ethnicity. Samples were projected onto the PC axes defined by the samples from 1000 Genomes Project (shown in light colors). **a)** The first two PCs can distinguish samples from three major continents. **b)** PC3 further distinguishes South Asians from Admixed Americans. Sample ethnicities were inferred based on the first four PCs using a machine learning approach implemented in peddy^56^.

**Table 1:** Phenotypic description of the SPARK pilot cohort.

**Supplementary Table 1:** All *de novo* SNV/indels in the affected offspring.

**Supplementary Table 2: A)** Burden of *de novo* variants in SPARK ASD trios and **B)** in published ASD trios. Likely gene disruptive (LGD) variants include frameshift indels, stop gain SNVs, and variants affecting canonical splice sites. Deleterious missense variants are defined by CADD score^10^ ≥ 25 or by MPC score^57^ ≥ 2. Genes are classified as constrained genes based on pLI≥0.5. The enrichment of observed de novo variants were compared to the baseline expectations^9^ by one-sided Poisson test. Baseline mutation rates were recalibrated so that the observed number of de novo silent mutations matches the expectation.

**Supplementary Table 3**: All singleton LGD variants (transmitted or un-transmitted) in known ASD/NDD genes. Singleton variants are defined as appearing only once in the SPARK pilot cohort. Rare singletons variants are singletons with ExAC allele frequency (all populations) < 0.001. Private singleton variants are singletons that are also absent from 1000 genomes, ESP, and ExAC databases.

**Supplementary Table 4**: All rare, inherited CNVs in the affected offspring.

**Supplementary Table 5:** All rare, de novo CNVs in the affected offspring.

**Supplementary Figure 3:** Parallel calling approach for mosaic SNVs.

**Supplementary Figure 4:** CUMC Method development: FDR-based minimum N_alt_ threshold. Variants called by samtools are shown here. **a)** Theoretical FDR-based minimum alternate allele read depth (N_alt_) thresholds as a function of total read depth (N). Assuming that sequencing errors are independent and that errors occur with probability 0.005, with the probability of an allele-specific error being 0.005/3=0.00167, and given the total number of reads (*N*) supporting a variant site, we iterated over a range of possible *N*_alt_ values between 1 and 0.5*N and estimated the expected number of false positives due to sequencing error, exome-wide [*(1-Poisson(N_alt_, λ=N*(0.00167))* 3×10^7^*]. Assuming one coding *de novo* SNV per individual^58^ and that roughly 10% of *de novo* SNVs arise post-zygotically^20–22^, we estimate there to be 0.1 mosaic mutations per exome. Under this assumption, to constrain theoretical FDR (in terms of distinguishing low allele fraction sites from technical artifacts) to 10%, we allowed a maximum of 0.01 false positives per exome. We used this cutoff to identify an FDR-based minimum *N*_alt_ threshold for each site as a function of total site depth. The dashed line denotes the threshold at which the expected number of false positives exome-wide is 0.01. **b)** FDR-based minimum N_alt_ threshold applied to samtools calls. Variant calls are plotted using total read depth (DP) and alternate allele read depth (N_alt_). The red line marks the N_alt_ cutoff as a function of DP.

**Supplementary Figure 5:** CUMC Method development: mosaic candidate identification. Data shown are the consensus variant calls. **a)** Total read depth (DP) in relation to variant allele fraction (VAF). The blue line denotes the Beta-Binomial mean VAF and the red lines denote the 95% confidence interval. To calculate the posterior odds that a given variant arose post-zygotically, we first calculated a likelihood ratio (LR) using two models: M_0_: germline heterozygous variant, and M_1_: mosaic variant. Under our null model M_0_, we calculated the probability of observing N_alt_ from a beta-binomial distribution with site depth *N*, observed mean germline VAF *p*, and overdispersion parameter *θ*. Under our alternate model M_1_, we calculated the probability of observing *N*_*alt*_ from a beta-binomial distribution with site depth *N*, observed site VAF *p*=*N*_*alt*_/*N*, and overdispersion parameter *θ*. Finally, for each variant, we calculated LR by using the ratio of probabilities under each model and posterior odds by multiplying LR by our EM estimated prior mosaic fraction estimate. Sites with posterior odds greater than 10 were predicted mosaic (corresponding to 9.1% FDR). **b)** Expectation-Maximization (EM) decomposition of variant allele fraction (VAF) into germline and mosaic distributions. Blue and red lines denote smoothed density curves for each distribution. We used an expectation-maximization (EM) algorithm to jointly estimate the fraction of mosaics among apparent *de novo* mutations and the false discovery rate of candidate mosaics. This initial mosaic fraction estimate gives a prior probability of mosaicism independent of sequencing depth or variant caller and allows us to calculate, for each variant in our input set, the posterior odds that a given site is mosaic rather than germline.

**Supplementary Figure 6:** Characterization of high confidence union mosaic calls. **A)** Alternative allele depth in relation to FDR based threshold. All calls are above the FDR threshold for both a 0.1 or 0.2 events per exome expectation. **a)** Variant allele fraction distribution. The grey and red bars denote germline and mosaic variants, respectively. The red line denotes the estimated true number of mosaics at each VAF window adjusted for mosaic detection power. Detection power is estimated as a function of variant allele fraction and sample average sequencing depth. The dashed vertical line denotes 5% VAF, below which estimated detection power is extremely limited and likely to artificially inflate adjusted counts. **c)** Percentile ranked distribution of Krupp et al.^22^ logistic mosaic score, 0.518 was the applied threshold for OHSU pipeline. Scores are overall well distributed between overlapping and group specific calls. Three of the CUMC only calls were not scored as they were filtered out by the OHSU pipeline before scoring due to differences in segmental duplication annotation.

**Supplementary Figure 7:** Comparison of Mosaic Mutations Spectra and Signatures in SPARK and SSC. Mutational contexts and frequency were extracted and plotted using the R package *MuationalPatterns*^59^. **a)** Mutational spectrum of the six different possible substitutions for SSC and SPARK mosaic mutations. **b)** Mutational signature of the relative frequency of mutations (Y-axis) within trinucleotides (context) for SSC and SPARK mosaic mutations. Though there are fewer calls in SPARK due to the smaller cohort size, both SSC and SPARK show a strong correlation to the same Cancer Signatures which are indicative of endogenous and DNA mismatch repair mutational processes.

**Supplementary Figure 8:** IGV plots used in mosaic mutation visualization and review. **A)** Example mosaic candidate passing IGV review – SP0026933:chr16:3777957:G>A **B)** Example mosaic candidate failing IGV review – SP0010023:chr19:48305658:G>A

**Supplementary Table 6:** List of likely mosaic variants.

**Figure 1:** Meta-analysis using the TADA framework provides statistical support for 34 genes at a false discovery rate (FDR) of 0.2. Known ASD genes are defined as those with SFARIgene^60^ score ≤ 2 or implicated in a previous TADA meta-analysis (FDR<0.1)^8^; known NDD genes are those listed in the DDG2P database^16^. D-mis variants are defined by CADD score ≥ 25. A total of 34 genes with at least one de novo damaging variant observed in SPARK pilot trios achieve an FDR<0.2 after meta-analysis with published trios (total n=5,230). Fourteen genes have not been previously classified as known ASD or NDD genes. Nine of these genes are supported by additional evidence (Supplementary Table 10) and are considered newly emerging ASD risk genes.

**Supplementary Table 7**: Results from the TADA meta-analysis of *de novo* variants from published simplex ASD trios (n=4,773) and SPARK pilot trios (n=457). Only genes with *de novo* LGD or D-mis (defined by CADD>25) variants observed in SPARK pilot trios are shown.

**Supplementary Note:** Further details on the justification of using an FDR of 0.2 in the TADA meta-analysis.

**Supplementary Table 8:** TADA analysis on simulated data of 5,238 trios. The number of positive findings, true positives, and fraction of false positives (FR) at different FDR thresholds are averaged over 200 repeats.

**Table 2:** Variants in newly emerging ASD risk genes in published and SPARK trios and associated phenotypic information.

**Supplementary Table 9:** Independent support for the nine newly emerging ASD risk genes and potential ASD risk genes identified by our TADA meta-analysis. Known ASD genes are defined as SFARIgene^60^ score ≤ 2 or identified in a previous TADA meta-analysis (FDR<0.1)^8^). Known NDD genes were defined as those listed in the DDG2P database^16^. We also systematically evaluated constrained genes (pLI>0.5) in which we identified de novo LGD variants in the SPARK cohort for which there are previously identified LGD variants or CNVs in individuals with developmental disorders. We also evaluated constrained genes (pLI>0.5) disrupted by deletions that we identified in SPARK, which affect less than five constrained genes and overlap with previously published copy-number deletions. For each gene, we checked membership in the following gene sets that were previously associated with ASD: **FMRP targets**: genes whose mRNA translation in neurons is likely regulated by the FMR1 protein, based on bioinformatics prediction and regulatory sequence motifs^33^; **PSD**: post-synaptic density components based on human neocortex proteomics^34^; **Embryonic**: genes whose expression levels are high in post-mortem embryonic brains and then decrease after birth, based on BrainSpan expression data and computationally derived by Iossifov et al. 2014^3^; **M2,M3,M16,M13**: Gene co-expression modules that are enriched for known ASD genes from a previous analysis of Parikshak et al 2013^44^; **Brain specific expression**: genes specifically expressed in fetal or adult brain, defined as expression index for the fetal or adult brain greater than the median expression for the entire data-set and greater than twice the median expression of non-brain tissue; based on the Novartis Tissue Expression Atlas and previously compiled by Yuen et al. 2015^5^; **Brain high expression**: genes that have log2 (RPKM) >= 4.86 and at least 5 BrainSpan data points, compiled by Yuen et al. 2015^5^; **Transcript regulation**: GO:0006355; **Chromatin modifier**: GO:0016569; **Nervous system development**: GO:0007399; **Nerve Impulse**: GO:0019227 (neuronal action potential propagation), GO:0019226 (transmission of nerve impulse), and GO:0050890 (cognition) and **Neuron projection**: GO:0043005. In addition, we also searched the literature for studies implicating the gene in central nervous system development. Genes were excluded from consideration if they were not supported by any line of evidence listed above.

**Supplementary Figure 9:** Support for newly emerging ASD genes from forecASD. The emerging genes highlighted in this work have significantly elevated forecASD scores (P=8.8×10^−7^, Wilcoxon test), with all genes in the top decile. Two constituent features in the forecASD ensemble (brain spatiotemporal expression and network topology) also show significantly elevated scores (P=0.017 and P=0.004, respectively), suggesting that these genes show similar properties to known ASD genes beyond genetic association and across a diverse feature space, thereby supporting the robust biological plausibility of these genes.

**Figure 2**: Network analysis and gene expression of newly emerging ASD genes using **A)** GO term clustering, **B)** PPI networks and **C)** gene expression of human fetal cortex at PCW (post-coital weeks) 15-16. Known ASD genes are defined as those with a SFARIgene score^60^ ≤2 (84 genes, indicated as SFARI) or implicated in a previous TADA meta-analysis^8^ at FDR<0.1 (65 genes, indicated as TADA). The enrichment for each gene was measured by the t-statistics comparing the expression level in each layer against all other layers. The enrichment of a gene set is the mean of t-statistics of its genes. Two newly emerging ASD risk genes (PAX5 and KDM1B) are not shown due to the low expression levels in human developing cortex (RPKM<1 for at least 20% available neocortical samples in BrainSpan^47^). Data were extracted from Supplementary Table of ^44^. Laminae abbreviations: marginal zone (MZ), outer/inner cortical plate (CPo/CPi), subplate (SP), intermediate zone (IZ), outer/inner subventricular zone (SZo/SZi), ventricular zone (VZ).

**Supplementary Table 10:** Summary of functional enrichment of network clusters depicted in Figure 2A.

**Supplementary Figure 10:** Gene expression of newly emerging ASD genes in human fetal brain PCW21.

**Supplementary Figure 11**: Expression specificity of newly emerging ASD genes in single-cell RNA-seq data from fetal and adult mouse brains. The specificity of expression in a cell type is measured by a specificity index which is the mean expression level in one cell type over the summation of mean expression level across all cell types^61^. For a gene set, the mean expression specificity of its genes was compared with 10,000 sets of randomly drawn genes matched for the transcript length and GC content and the enrichment is measured by the standard deviation from the mean specificity of random gene sets^61^. The mouse neuronal cell types are defined by the analysis of single cell RNA-seq data of fetal and adult mouse brains generated by Karolinska Institutet (KI) and used in the previous study^48^. The mouse orthologs of human genes were retrieved from MGI database^62^. The known ASD genes show highest enrichment in pyramidal neurons (in hippocampus CA1 and somatosensory cortex), cortical interneurons, and medium spiny neurons. The first three enriched cell types were previously reported for the 65 autism genes identified from TADA meta-analysis^61^. The newly implicated genes also show highest specificity in pyramidal neurons, suggesting functional convergence in these cell types.

**Supplementary Figure 12** Expression specificity of newly emerging ASD genes in single-cell RNA-seq data from human brains. Human neuronal cell types are defined by the single-nucleus RNA-seq data of archived human brains^49^. Known and new ASD genes were mostly enriched in neurons (exCA, exDG, exPFC) and interneurons (GABA). Highest enrichment was also observed in pyramidal neurons (exXCA). New ASD genes were also enriched in neuronal stem cells that are not implicated by known ASD genes, but the enrichment is not significant. Significance code: *= p<0.01, **= p<0.001. exPFC=glutamatergic neurons from the PFC, exCA=pyramidal neurons from the hippocampus CA region, GABA=GABAergic interneurons, exDG=granule neurons from the Hip dentate gyrus region, ASC=astrocytes, NSC=neuronal stem cells, MG=microglia, ODC=oligodendrocytes, OPC=oligodendrocyte precursor cells, NSC=neuronal stem cells, SMC=smooth muscle cells, END=endothelial cells.

**Supplementary Table 11**: All pathogenic (returnable) and possibly ASD-associated genetic variants.

**Supplementary Figure 13:** Eight pedigrees with multiple contributing variants.

**Supplementary Figure 14:** Genetic causes of ASD were identified in 6 offspring in 5 multiplex families, 3 of which are shown in this figure. SF0003496 and SF0008074 are shown in Supplementary Figure 13. SF0003496 has another affected offspring, who was not sequenced in this study.

**Supplementary Table 12**: Statistical power of TADA analysis. To simulate mutation data across all protein coding genes, we first randomly assigned each gene as ASD gene with probability of 0.05. Then for each ASD gene, we sampled relative risk (RR_i_) for LGD and D-mis variants from prior distributions Gamma(18,1) and Gamma(6,1) which were the same as used in TADA analysis; for non-ASD gene, relative risk will be 1 for both types of variants. Then the number of observed de novo variants of class c for gene i will be sampled from Poission(2*N*u_i,c_*RRi), where u_i,c_ is the baseline mutation rate and RRi is the relative risk. After generating the full data from all genes, we applied TADA to the dataset, and the procedure was repeated 100 times for each sample size. The table shows the average number of total positive findings and true positives at different FDR thresholds.

**Supplementary Figure 15**: Recalibrating VQS LOD threshold for analyzing inherited singleton variants. The transmission to un-transmission ratio of singleton synonymous SNVs **(A)** and non-frameshift indels **(B)** are shown as a function of the VQS LOD score. The dashed lines mark the GATK defined cutoffs based on different tranche sensitivity thresholds. The red line shows the cutoffs that balance the transmission to non-transmission ratio and were used in filtering singleton variants for transmission disequilibrium analysis.

### Data and code availability

The genomic and phenotypic data for the 1379 individuals analyzed in this study is available by request from SFARIBase (https://www.sfari.org/resource/sfari-base/) with accession ID: SFARI_SPARK_WES_p. Methods for SNV, Indels, CNV analysis are available at https://genomicpipelines.sparkforautism.org/

## Methods

### Participant recruitment, phenotyping and DNA sequencing

All participants were recruited to SPARK under a centralized IRB protocol (Western IRB Protocol #20151664). Participants are asked to fill out questionnaires online as described here: https://www.sfari.org/spark-phenotypic-measures/. Families are classified as multiplex if the initial individual with ASD registered in the study has a first-degree family member with ASD, as indicated either by enrollment or survey report. When possible, phenotypic severity was systematically assessed in all individuals in SPARK with a genetic finding related to ASD and all individuals in SPARK and SSC with a likely deleterious genetic variant in any of the nine newly implicated ASD risk genes.

In SSC and SPARK participants, severity ratings were assigned as follows. Individuals were given 0 to 3 points based on the level of intellectual disability (ID) as indicated by IQ scores (SSC) or parent report (rated 0=no ID, 1=mild ID, 2=moderate ID, 3=severe ID). ID was rated based upon all available information related to degree of impairment or disability in functioning (such as encopresis or nonverbal status in an older child or adult). Individuals were given additional severity points if there was a history of seizures (1 point) or if birth defects were present (1 point). Macrocephaly was not included as a birth defect due to its prevalence in ASD. ID is weighted by a factor of two, for its impact on functioning. In the absence of clear ID, a developmental delay was rated as 1, unweighted.

Essential phenotypic information was curated across language and motor development, co-morbidities, and Repetitive Behavior Scale-Revised^63^, Social Communication Questionnaire-Lifetime^64^ and Developmental Coordination Disorder Questionnaire score^65^ (Table 2). In SSC, all phenotype details were determined through clinic evaluation and interview; specifically, language delay was defined by Autism Diagnostic Observation Schedule module (1 through 4) per age^66^, and regression was determined from the Autism Diagnostic Interview-Revised^67^. For SPARK, all variables were taken from parent report. It was noted that rates of language disorder and psychiatric co-morbidities are lower in SSC likely due to DSM-IV diagnostic practice at the time.

Saliva was collected using the OGD-500 kit (DNA Genotek). Exome capture was performed using VCRome and the spike-in probe set PKv2. Captured exome libraries were sequenced using the Illumina HiSeq platform in 100 bp paired end reads. The Illumina HumanCoreExome (550K SNP sites) array was used for genotyping.

### Read alignment and QC

Post-sequencing reads were aligned to build 37 of the human genome using bwa version 0.6.2-r126^68^, duplicates were marked using Picard version 1.93 MarkDuplicates, and indels were realigned using GATK^69^ version 2.5-2-gf57256b IndelRealigner. Quality checks were performed on the BAM files using SAMTools^70^ version 1.3.1 flagstat and Picard version 2.5.0 CalculateHsMetrics. Overall, 98 ± 1.8% of the reads mapped to the genome, 96 ± 2.3% of the reads were properly paired reads, and 87 ± 15% of targeted regions had >=10X coverage.

KING^71^ was used for relatedness inference based on the genotype of exome SNPs (MAF>0.01). Estimated kinship coefficient and number of SNPs with zero shared alleles (IBS0) between a pair of individuals were plotted. Parent-offspring, sibling pairs, and unrelated pairs can be distinguished as separate clusters on the scatterplot (Supplementary Fig 1). One outlier parent-offspring pair (SP0002452 and mother) showed higher than expected IBS0 and was caused by parental chr6 iso-UPD. Pairwise scatterplots of heterozygotes to homozygotes (het/hom) ratio of chromosome X, sequencing depth of chromosome X and Y normalized by the mean depth of autosomes were used for sex check. Two samples with sex chromosome aneuploidy were identified as outliers in the scatterplot (Supplementary Fig. 2).

### Variant calling

#### *De novo* SNV/indel detection

*De novo* sequence variants were called by three groups - University of Washington (UW), Simons Foundation (SF), Columbia University Medical Center (CUMC) - according to the methods below.

#### UW

Variants were called from whole exome sequence (WES) using FreeBayes^72^ and GATK^69^. FreeBayes version v1.1.0-3-g961e5f3 was used with the following parameters: --use-best-n-alleles 4 -C 2 -m 20 -q 20; and GATK version 3.7 HaplotypeCaller was used with the following parameters: -A AlleleBalanceBySample -A DepthPerAlleleBySample -A MappingQualityZeroBySample -A StrandBiasBySample -A Coverage -A FisherStrand -A HaplotypeScore -A MappingQualityRankSumTest -A MappingQualityZero -A QualByDepth - A RMSMappingQuality -A ReadPosRankSumTest -A VariantType. Post-calling bcftools ^73^ version 1.3.1 norm was used with the following parameters -c e -O z -s -m –both. We identified candidate *de novo* calls based on the intersection of FreeBayes and GATK VCF files and identifying variants present in offspring but not in parents. We required a minimum of 10 sequence reads in all members of the parent-offspring trio; an allele balance > 0.25 and a PHRED quality >20 for both FreeBayes and GATK variants.

#### SF

Sequence data were preprocessed using GATK best practices and variant calls were predicted using three variant callers: GATK v3.6^74^, FreeBayes v1.1.0-441 and Platypus v0.8.1-0^75^. GATK: gVCF files were generated for each sample with GATK HaplotypeCaller (minimum confidence thresholds for calling and emitting was set to 30 and 10, respectively); joint variant calls were performed using GATK GenotypeGVCFs with the recommended default hard filters. For SNPs, we filtered out: QD < 2.0 ‖ FS > 60.0 ‖ MQ < 40.0 ‖ MQRankSum < −12.5 ‖ ReadPosRankSum < −8.0. For indels, we filtered out: QD < 2.0 ‖ FS > 200 ‖ ReadPosRankSum < −20.0. FreeBayes: variants were called with default settings for optimal genotyping of indels in lower-complexity sequence. The final data set included candidate calls with a quality of 5 or greater. Platypus variant calling was performed with local assembly analysis when at most ten haplotypes were allowed. Variants were filtered out for allele bias (p-value < 0.0001), bad reads (>0.9), sequence complexity (>0.99) and RMSMQ (<20); other filters were applied on estimated haplotype population frequency (FR), total coverage at the locus (TC) and phred-scaled quality of reference allele (QUAL): (FR[0] <= 0.5 and TC < 4 and QUAL < 20),or (TC < 13 and QUAL < 10),or (FR[0] > 0.5 and TC < 4 and QUAL < 50). For each variant caller, a variant was identified as a candidate *de novo* variant if the variant was called in the proband and it occurred only once in the cohort, with an alternative allele fraction between 0.2 and 0.8. Both parents were required to have the homozygous reference genotype at the *de novo* locus. Read coverage of the variant locus had to be at least ten reads in each sample in the trio. *De novo* candidate variants were classified by DNMFilter algorithm^76^ that was re-trained with the SSC data set^3,14^: 1800 de novo mutations identified by both Iossifov et al, 2014^3^ and Krumm et al, 2015^14^, 1104 validated SNVs and indels from both studies and 400 variants that failed validation. We also randomly selected ~ 3000 negative examples from the pool of all SSC variants that were not confirmed to be de novo. After merging *de novo* candidate variants from three variant callers, candidate *de novo*s were considered if they occurred only once in the cohort, passed hard filters, and had assigned *de novo* probability greater than 0.88 for SNVs and greater than 0.0045 for small indels. In the latter case, the total parental alternative allele count < 3 reads.

#### CUMC

Variants were called from aligned sequence data using GATK HaplotyperCaller to generate individual level gVCF files. All samples in the cohort were then jointly genotyped and have variant quality recalibrated by GATK v3.8^69^. A variant present in the offspring with homozygous reference genotypes in both parents was considered to be a potential *de novo* variant. We used a series of filters to identify *de novo* variants. Briefly, we included variants that passed VQSR filter (tranche≤99.7 for SNVs and ≤99.0 for indels) and had GATK’s Fisher Strand≤25, quality by depth≥2. We required the candidate *de novo* variants in probands to have ≥5 reads supporting alternative allele, ≥20% alternative allele fraction, Phred-scaled genotype likelihood ≥60 (GQ), and population allele frequency ≤0.1% in ExAC; and required both parents to have >=10 reference reads, <5% alternative allele fraction, and GQ≥30.

#### *De novo* SNV/indel consensus call set and annotation

*De novo* variants were independently called by three centers – UW, SF, CUMC. *De novo* variants called by all three groups were included in the final list by default. Those called by one or two groups were manually evaluated and included in the final list if consensus is reached among all groups after discussion and manual inspection with IGV plots. Variants were annotated by ANNOVAR^77^ based on GENCODE Basic v19^78^. Candidate mutations in the ACMG secondary findings v2 59 gene list^79^ (except *PTEN*, *TSC1*, *TSC2* and *NF2*, which are genes included in the SPARK ASD genes list) were excluded. Coding *de novo* variants - nonsense, missense, or synonymous single nucleotide variants (SNVs), frameshift or non-frameshift indels, and splicing site variants - were annotated. *De novo* variants were also annotated with snpEff version 4.1g^80^ (reference GRCh37.75), SFARIgene scores (version q1, 2018, https://gene.sfari.org/database/gene-scoring/), CADD^10^, MPC^11^ and findings from Deciphering Developmental Disorders project (gene2phenotype).

#### Inherited singleton variants

We first performed following filtering on individual genotypes. We required minimal read-depth >=10 and GQ>=30, required allelic balance <0.1 for homozygotes reference, >0.9 for homozygotes alternative, and 0.3~0.7 for heterozygotes SNVs (0.25~0.75 for heterozygotes indels). Genotype calls not passing those criteria were set to missing. Then we removed variants having missing genotypes in >25% of founders. We focused analysis on singleton variants in which the alternative allele was only seen in one parent in the data. We calibrated GATK’s VQS LOD score for SNV and indels separately such that synonymous singleton SNVs and non-frameshift singleton indels were transmitted 50% of the time (Supplementary Figure 14) The resulting VQS LOD score cutoffs are −1.85 for SNVs and −1.51 for indels. As mentioned in the Result section, inherited LGD variants are less likely cause complete loss of function to the gene. To prioritize inherited LGD variants, we require the variant to be annotated as HC (high confidence) by LOFTEE v0.3^12^ using default parameters in >60% of the GENCODE transcripts.

#### Identification of Mosaic Mutations

Mosaic SNVs were independently called by two centers – Oregon Health & Science University (OHSU) and CUMC. The OHSU approach was previously published^22^ and utilized a binomial deviation and logistic regression model to score candidate mosaic variants. The CUMC approach used a novel approach that was based on a beta-binomial deviation and an FDR based approach to determine per site thresholds.

#### OHSU

SNVs were called as previously described^22^. In brief, pileups were generated using SAMtools (v 1.1) with BAQ disabled and mapQ 29 (*samtools mpileup –B –q 29 –d 1500*) on processed BAMs. Variants were called on individual samples using VarScan 2.3.2, LoFreq 2.1.1 and an in-house mpileup parsing script (mPUP). Additional parameters for Varscan included: --*min-var-freq 1×10^−15^ –p-value 0.1*. Per sample caller outputs were combined and annotated using ANNOVAR (03/22/15 release) with databases: Refseq genes (obtained 03/2017), segmental duplications (UCSC track genomicSuperDups, obtained 03/25/2015), repetitive regions (UCSC track simpleRepeat and hg19_rmsk, obtained 03/25/2015), Exome Aggregation Consortium (ExAC) release 0.3 (obtained 11/29/2015), Exome Sequencing Project (ESP) 6500 (obtained 12/22/2014), and 1000 Genomes Phase 3 version 5 (obtained 12-16-2014).

Variants were filtered based on the best practices established in Krupp et al. 2017^22^: 1) variant must be exonic or disrupt a canonical splice site, 2) have a population frequency of <= 0.5%, 3) have at least five alternative reads, 4) not be in a known segmental duplication or repetitive regions (SDTRF), 5) called by at least two variant callers, 6) SPARK cohort count <= 1 and SSC cohort count <= 2, 7) variant read mismatch <= 3, and 8) allele fraction upper 90% confidence interval >= 0.05. For a variant to be considered *de novo*, parental alternative allele count must be <= 4 reads. *De novo* variants were considered to be candidate mosaic variants if: the probability the allele fraction significantly deviated from heterozygous (PHET) was <= 0.001, 2) the allele fraction upper 90% confidence interval was < 0.4, and 3) a logistic regression model score was >= 0.518.

#### CUMC

SNVs were called on a per-trio basis using SAMtools (v1.3.1-42) and BCFtools (v1.3.1-174). We generated trio VCF files using samtools ‘*mpileup’* command with options *‘–q 20 –Q 13*’ corresponding to mapQ and baseQ thresholds of 20 and 13 respectively, followed by bcftools ‘*call’* with option ‘*–p 1.1*’ to expand the set of variant positions to be evaluated for mosaicism. In contrast to the OHSU pipeline, BAQ was used to potentially reduce false positive SNV calls caused by misalignments^81^. To identify *de novo* variants from trio VCF files, we selected for sites with (i) a minimum of six reads supporting the alternate allele in the proband and (ii) for parents, a minimum depth of ten reads and 0 alternate allele read support. Variants were then annotated using ANNOVAR (v2017-07-17) to include information from refGene, gnomAD (March 2017), 1000 Genomes (August 2015), ExAC, genomicSuperDups, COSMIC (v70), and dbSNP (v147) databases. CADD^10^, MPC^11^ were used to annotate variant functional consequence.

##### Pre-processing and QC

To reduce the noise introduced by our variant calling approach, we preprocessed our variants using a set of filters. Since our method is allelic depth-dependent, we took a conservative filtering approach to reduce the impact of false positives on model parameter estimation. We first filtered our variant call set for rare heterozygous coding variants (MAF<=1×10^−4^ across all populations represented in gnomAD and ExAC databases). To account for regions in the reference genome that are more challenging to resolve, we removed variant sites found in regions of non-unique mappability (score<1; 300bp), likely segmental duplication (score>0.95), and known low-complexity^82^. We then excluded sites located in *MUC* and *HLA* genes and imposed a maximum variant read depth threshold of 500. To account for common technical artifacts, we used SAMtools PV4 p-values with a threshold of 1×10^−3^ to exclude sites with evidence of baseQ bias, mapQ bias, and tail distance bias. To account for potential strand bias, we used an in-house script to flag sites that have either (1) 0 alternate allele read support on either the forward or reverse strand or (2) p<1×10^−3^ and OR<0.33 or OR>3 when applying Fisher’s method to compare strand based reference or alternative allele counts. Finally, we excluded sites with frequency >1% in the SPARK pilot, as well as sites belonging to outlier samples (with abnormally high *de novo* SNV counts, cutoff = 7) and complex variants (defined as sites with neighboring *de novo* SNVs within 10bp).

##### IGV Visualization of Low Allele Fraction de novo SNVs

To identify likely false positives among our low allele fraction (VAF<0.3) *de novo* SNVs, we used Integrative Genomics Viewer (IGV v2.3.97) to visualize the local read pileup at each variant across all members of a given trio. We focused on the allele fraction range 0.0-0.3 since this range captures the majority of the technical artifacts that will negatively impact downstream parameter estimation. Sites were filtered out if (1) there were inconsistent mismatches in the reads supporting the mosaic allele, (2) the site overlapped or was adjacent to an indel, (3) the site had low MAPQ or was not primary alignment, (4) there was evidence of technical bias (strand, read position, tail distance), or (5) the site was mainly supported by soft-clipped reads.

##### Empirical Bayes Post-zygotic Mutation Detection Model

To distinguish variant sites that show evidence of mosaicism from germline heterozygous sites, we modeled the number of reads supporting the variant allele (*N*_*alt*_) as a function of the total site depth (*N*). In the typical case, *N*_*alt*_ follows a binomial model with parameters *N* = site depth and *p* = mean VAF. However, we observed notable overdispersion^83,84^ in the distribution of variant allele fraction compared to the expectations under this binomial model. To account for this overdispersion, we instead modeled *N*_*alt*_ using a beta-binomial distribution. We estimated an overdispersion parameter *θ* for our model whereby for site depth values *N* in the range 1 to 500, we (1) bin variants by identifying all sites with depth *N*, (2) calculate a maximum-likelihood estimate *θ* value using *N* and all *N*_*alt*_ values for variants in a given bin, and (3) estimate a global *θ* value by taking the average of *θ* values across all bins, weighted by the number of variants in each bin.

We used an expectation-maximization (EM) algorithm to jointly estimate the fraction of mosaics among apparent *de novo* mutations and the false discovery rate of candidate mosaics. This initial mosaic fraction estimate gives a prior probability of mosaicism independent of sequencing depth or variant caller and allows us to calculate, for each variant in our input set, the posterior odds that a given site is mosaic rather than germline.

#### Finalized Union Mosaic Call Set and Validation Selection

The high confidence call sets from the two parallel mosaic determination approaches were combined, and all candidate mosaic variants were then inspected manually in IGV. Variants in regions with multiple mismatches or poor mapping quality were removed, and the remaining mosaics comprised the high confidence mosaic call set. For calls that were unique to one approach, the variant was annotated with which quality filter it initially failed. Variants that were flagged as low confidence germline by CUMC approach but mosaic by OHSU approach had posterior odds > 1 and were thus retained in the union call set.

#### CNV detection

*De novo* and rare inherited CNVs were independently called by two centers - UW and SF. The final CNV list included all autosomal CNVs that were called by both SF and UW pipelines either with reciprocal overlap of at least 50% or when the CNV from one pipeline was completely within the CNV from the other pipeline. In both cases, the overlapping region was reported as the final region and annotated as described below. CNVs called only by one pipeline were considered as high confidence CNVs if they were called by at least 2 tools or if they were de novo CNVs confirmed by manual inspection of plots on exome data. High confidence CNVs were also included in the final list after discussion and manual inspection of plots on exome data. De novo CNVs were additionally inspected on BAF and LRR plots on genotyping data. CNVs that had at least 75% overlap with known segmental duplications (segDups track for hg19 from UCSC browser) were excluded. All CNVs were annotated with the list of RefSeq HG19 genes, OMIM genes, brain embryonically expressed genes^3^, brain critical genes^18^, ASD significant^85^ and ASD related genes^8,14^ that have their coding regions overlapping with the CNV. In addition, each found gene was annotated with pLI (ExAC release 0.3, http://exac.broadinstitute.org/downloads), ASD^86^, RVIS^87^, LGD^87^, and SFARIgene scores (version q1, 2018, https://gene.sfari.org/database/gene-scoring/). De novo CNVs that affect DUSP22 and olfactory genes were excluded due to high variability in copy-number of those regions among individuals^88^.

#### UW, detection using XHMM and CoNIFER

CNVs from WES were called using CoNIFER and^89^ XHMM^90^. CoNIFER version v0.2.2 was used with the S value, --svd 7, set as a threshold as suggested by the scree plot. XHMM version statgen-xhmm-3c57d886bc96 was used with the following parameters --minTargetSize 10 --maxTargetSize 10000 --minMeanTargetRD 10 --maxMeanTargetRD 500 --minMeanSampleRD 25 --maxMeanSampleRD 200 -- maxSdSampleRD 150 to filter samples and targets, and then to mean-center the targets; PVE_mean --PVE_mean_factor 0.7 was used to normalize mean-centered data using PCA information; --maxSdTargetRD 30 was used to filter and z-score centers (by sample) the PCA normalized data; and then to discover CNVs in all samples. Calls from CoNIFER and XHMM were merged in a VCF file using https://github.com/zeeev/mergeSVcallers with the following parameters -t xhmm,conifer -r 0.5 -s 50000, then merged VCF was sorted by Picard version v2.5.0, and zipped and indexed with Tabix version v0.2.6. We re-genotyped each XHMM and CoNIFER CNV event by assessing the RPKM values from the CoNIFER workflow on an individual. Probands were considered to have a deletion if their average RPKM value was less than −1.5 s.d and have a duplication if their average RPKM value was greater than 1.5 s.d. For an event to be considered as variant in a parent, we required an average ZRPKM less than −1.3 or greater than 1.3 for deletions and duplications, respectively.

#### UW, CNV validation using SNP microarray

We generated an independent CNV callset for validation purpose using SNP microarray genotyping data generated from Illumina InfiniumCoreExome-24_v1.1, where IDATs (n=1,421) were processed using Illumina Genome Studio Software. CNV analysis was performed using the Illumina CNVpartition algorithm version v3.2.0. Log R Ratio data for all samples and probes was exported. PennCNV^91^ version v1.0.4 was used to detect CNVs with the following parameters -test –hmm -pfb all.pfb --gcmodelfile –confidence. We determined the maximum and minimum overlap of SNP microarray CNVs based on the presence of WES probes to make the array calls more similar to the exome calls and considered an event to have support by PennCNV or CNVpartition if there was at least 50% reciprocal overlap. We also generated per probe copy number estimates using CRLMM^92,93^ version 1.38.0 as previously described^14^ and genotyped each candidate WES CNV. Deletions were considered variant if they had a p-value less than 0.05 and a mean percentile rank less than 30. Duplications were considered variant if they had a p-value less than 0.05 and a percentile rank of mean greater than 70. CNVs passing the RPKM genotyping were combined with the CNV data from CRLMM, PennCNV, and CNVPartition. We considered WES CNVs as valid if there was support for gain or loss from the PennCNV, CNVpartition, or CRLMM approaches described above. We assessed inheritance using both SNP and WES data and preferentially scored inherited events over *de novo* CNVs.

#### SF

CNVs were called with two tools - xHMM v 1.0^94^ and CLAMMS v 1.1^95^. xHMM: CNVs were called with default settings (except not filtering on the maximum target size), including filtering low complexity and GC extreme targets. CLAMMS: CNVs were called with INSERT_SIZE=390 bp and training per-sample-models on sample specific reference panels due to the observed batch effect in the data; CLAMMS calls were filtered for all CNVs with Q_EXACT less than 0, or Q_SOME less than 100, or CNVs that were in samples with more than 70 predicted CNVs of the size at least 10 Kb and of quality score Q_SCORE at least 300. The inheritance status of the autosomal CNVs was determined by default xHMM protocol for *de novo* CNVs identification with plink 1.07^96^ and Plink/Seq 0.10 [https://atgu.mgh.harvard.edu/plinkseq/]. Similar protocol was implemented in java for CLAMMS analysis. For each tool, two tiers of CNV calls– the most confident calls (tier 1) and less confident calls (tier 2) - were defined, based on *de novo* and transmission rates for different cuts on quality scores: SQ (phred-scaled quality of some CNV event in the interval) and NQ (phred-scaled quality of not being diploid, i.e., DEL or DUP event in the interval) in xHMM and Q_SOME (phred-scaled quality of any CNV being in this interval) in CLAMMS. xHMM tier1 included all autosomal CNVs with both SQ and NQ quality scores of at least 60, and tier2 - all autosomal CNV calls with quality scores between 30 and 60. Samples with more than 10 *de novo* CNVs in xHMM tier1 of size at least 10 kb were excluded. CLAMMS tier1 included all predictions with quality score 999, except predictions for 25 probands that have CNVs of size greater than 500 kb with quality score 999 or predictions, which region was partially inherited and partially *de novo*; tier 2 included those excluded from tier 1 predictions as well as all CNVs with quality score Q_SOME at least 400 and less than 999. Predictions by both methods that had less than 3 exons or at least 75% overlap with known segmental duplications (segDups track for hg19 from UCSC browser) were removed from the list. The final list of CNV predictions included all CNVs from tier 1 predicted by either xHMM or CLAMMS and “intersection” of tier 2 sets from both tools, that is, CNVs that were confirmed by two tools with reciprocal or cumulative reciprocal overlap of at least 50%. In the latter case, CNV predicted by one tool is covered by a set of CNVs predicted by the other tool. If a CNV from xHMM or CLAMMS was confirmed by the other tool, the overlapping region was reported as the final region. CNVs were removed from the analysis if it had more than half of its length overlapping with the ACMG secondary findings v2 gene^79^ (except *PTEN, TSC1, TSC2* and *NF1* genes in the ASD genes list). If such gene covers less than 50% of CNV, the part of CNV without the gene was kept if it has at least 25% of its length not covered by segmental duplications. To identify higher confidence CNV predictions, xHMM and CLAMMS plots were manually investigated for each CNV in the final SF list. In addition, SF predictions were compared with PennCNV^91,97,98^ calls from array data, which have confidence score of at least 100. All reciprocal overlaps of at least 50% were treated as additional evidence for CNV support.

#### UW, chromosome aneuploidy assessment

We also assessed evidence of chromosomal aneuploidy by calculating sequence read depth using SAMTools^10^ version 1.4 on a per chromosome basis normalizing by the relative density of WES probes and comparing the normalized value for each chromosome to the normalized value on chromosome 1 (assumed to be diploid). For autosomes, we multiplied this number by two to get the estimate of chromosomal copy number. We did not multiply by two for the X or Y chromosomes. To further assess the chromosomal copy number, the heterozygosity was calculated for all SNPs and indels. For heterozygous sites, the absolute mean deviation from 0.5 was also calculated. We assessed both metrics to identify outliers. Aneuploidies were required to have support from both the read depth and SNP/indel metrics.

### Burden of *de novo* variants

Baseline mutation rates for different classes of *de novo* variants in each GENCODE coding gene were calculated using a previously described mutation model^9^. Briefly, the trinucleotide sequencec context was used to determine the probability of each base mutating to each other possible base. Then, the mutation rate of each functional class of point mutations in a gene was calculated by adding up the mutation rate of each nucleotide in the longest transcript. The rate of frameshift indels was presumed to be 1.1 times the rate of nonsense point mutations. The expected number of variants in different gene sets were calculated by summing up the class-specific variant rate in each gene in the gene set multiplied by twice the number of patients (and if on chromosome X, further adjusted for female-to-male ratio^99^).

The observed number of variants in each gene set and case group was then compared with the baseline expectation using a Poisson test. In all analyses, constrained genes were defined by a pLI score of ≥0.5. To compare with previously published ASD studies, we collected published *de novo* variants identified in 4773 simplex trios from three largest autism studies to date^3,4,7^. To account for platform differences, the baseline mutation rate of each gene was scaled so that the exome-wide expected number of silent variants matches the observed count.

### TADA analysis

To perform TADA analysis of de novo variants, we assumed the fraction of disease genes is 5% as estimated by previous studies^25,100^. The prior relative risk for LGD variants and D-mis (defined by CADD>=25) were specified as Gamma (18,1) and Gamma (6,1). The prior mean relative risks were determined using the relationship between burden and relative risk as described previously^25^. The baseline mutation rate of each gene was the same as used in burden analysis. The analysis was performed on de novo variants of 4773 published trios and after combing de novo variants identified from SPARK pilot trios.

### Laminal layer and cell type enrichment

To evaluate the expression specificity of laminal layer of human developing cortex, we analyzed RNA-seq data of neocortical samples of BrainSpan^47^ following the method of Parikshak et al 2013^44^. The expression specificity was measured by t-statistic comparing the expression level in each layer against all other layers. Two newly emerging genes (*PAX5, KDM1B*) were not included in the analysis due to the low expression levels (RPKM<1 for at least 20% available neocortical samples). To evaluate cell-type specificity, we used a published data of mouse neuronal cell types inferred from analyzing single cell RNA-seq data of fetal and adult mouse brains generated by Karolinska Institutet (KI)^101^, and human CNS cell types inferred from a single nucleus RNA-seq data^49^. The mouse orthologs of human genes were retrieved from MGI database^62^. The cell-type specificity was measured by a specificity index which is the mean expression level in one cell type over the summation of mean expression level across all cell types^61^. To analyze the overall trend of specificity of a gene set, the mean specificity measure of its genes was compared with 10,000 sets of randomly drawn genes matched for the transcript length and GC content and the enrichment is measured by the standard deviation from the mean specificity of random gene sets^61^.

### Network and functional analysis

The network depicted in Figure 2A was constructed using the top decile of forecASD genes, SFARI genes scoring 1 or 2, and SPARK newly implicated genes (9 total). These genes were projected onto the STRING network^102^ (v10) using the igraph R package (1708 genes). Edges within the STRING network were thresholded at 0.4, according to the authors’ recommendation. The largest connected subcomponent (1664 genes) was then extracted as the basis for further network analysis. Clustering was performed on the fully connected network using the fastgreedy.community function available within the igraph package. Clusters with fewer than 30 genes were not considered for further analysis (none of these clusters contained the nine genes highlighted here). Following the first round of clustering, clusters with >150 genes were subject to an additional round of clustering, with the goal of separating broad functions of genes into more specific subcomponents. This process resulted in 11 clusters. Each cluster was assessed for functional enrichment using the Gene Ontology^103^ as accessed through the clusterProfiler package within R. During the functional analysis the background gene universe was always set to the full set of genes represented among the 11 clusters. Visualization of this network analysis was performed in Cytoscape^104^. The top 5 most significant GO terms associated with each cluster are available in the Supplementary Table 10. Cluster labels in Figure 2 were chosen as the most representative among the top terms for each cluster. Figure 2B was constructed using the subset of the larger network (Figure 2A), corresponding to SPARK newly implicated genes and SFARI genes scoring 1 or 2 (88 genes). These genes were projected onto the STRING network within Cytoscape using the STRINGapp. All nonzero-weighted edges were considered. The fully connected component was visualized, which resulted in two genes being dropped (DEAF1 and RANBP17). Edges adjacent to newly-implicated genes with a STRING interaction score of ≥ 0.4 are highlighted.

#### ForecASD Analysis

We used a recently developed method, forecASD^38^ that indexes support for a gene being related to autism by integrating genetic, expression, and network evidence through machine learning. We examined the forecASD scores of the nine newly emerging ASD genes from the TADA analysis and compared them to the remainder of the genome using a Wilcoxon rank-sum test. We similarly used the Wilcoxon test and employed two predictive features used by forecASD (BrainSpan_score and STRING_score) to assess whether the new genes showed similarity to known ASD risk genes in terms of brain expression patterns and network connectivity. Importantly, because forecASD uses previously published TADA scores among its predictive features, which are strongly correlated with updated TADA scores, we investigated whether the elevated forecASD scores in our candidate genes could be explained solely by the previous TADA scores. Specifically, we fit a logistic regression model with the nine newly emerging genes labeled as ‘1’ and 500 size-matched background genes (not listed in the SFARI gene database) labeled as ‘0’ in the dependent variable (Y). Separate models were fit using either forecASD or TADA^8^ scores as predictors, or both together in a full model. Both TADA and forecASD were significantly associated with the “new gene” indicator when considered in isolation (P<0.001 for both). However, when included together in a model of Y, forecASD remained significantly associated (P=0.00012) while TADA lost significance (P=0.41). The Akaike information criterion (AIC) indicated that the forecASD-only model was a more optimal fit compared to either the TADA-only or TADA+forecASD fit. This analysis suggests that the elevated forecASD scores observed in the nine new genes cannot be fully explained by the use of TADA as a predictor in forecASD.

### Supplementary Note 1: FDR control in TADA analysis

In our TADA analysis, we combined *de novo* variants of published trios with SPARK and only presented the results for genes with damaging *de novo* variants in SPARK and whose q-values fall below some false discovery rate (FDR) threshold. It is important to note that q-value and FDR threshold are defined with respect to genome-wide genes without conditioning on observing variants in SPARK, because the TADA framework estimates FDR by modeling the genome-wide “alternative hypothesis” (i.e., number of risk genes and relative risks for different types of variants). By thresholding on genome-wide FDR, the procedure may not control the FDR among genes with variants in SPARK at the same cutoff.

To understand the effect of conditioning on variants in a subset of samples on the FDR among selected genes, we carried out the following simulation using the parameters of genetic architecture as we used as priors in TADA analysis. First, across all protein coding genes, we randomly assigned each gene as an ASD gene with a probability of 0.05. Second, for each ASD gene, we sampled relative risk (RR_i_) for LGD and D-mis variants from prior distributions of Gamma(18,1) and Gamma(6,1), respectively; for non-ASD genes, relative risk will be 1 for both types of variants. Then the number of observed *de novo* variants of class c for each ASD gene will be sampled from Poission(2*N*u_i,c_*RR_i,c_) distribution, where u_i,c_ is the baseline mutation rate and RR_i,c_ is the relative risk. After generating *de novo* variants for all genes in a cohort of 5,238 (4,773+465) trios which is the same as combined published and SPARK trios, we applied TADA to the simulated data first on the subset of 4,773 trios and then on all the full cohort. The overall procedure was repeated 200 times. We found that by conditioning on observing variants in the subset of trios, the average FDR among selected genes are consistently lower than the specified threshold (Supplementary Note 1, Supplementary Table 8 left). For example, at the genome-wide FDR cutoff of 0.2, we observe on average 45 genes with variants in the 465 trios with the proportion of false positives of 15.9%. Intuitively, this is because true disease genes are more likely to have recurrent de novo variants in the new data.

We also considered a second scheme of selecting genes: by focusing on those that are initially not significant but fall below desired FDR threshold after the inclusion of new data (newly significant). We found that the average FDR among newly significant genes is typically higher than the specified threshold (Table 1, right). For example, by the same simulation and focus on genes with q-value>0.2 in analyzing 4773 trios but <=0.2 after inclusion of additional 465 trios, we observe on average 25 newly significant genes with the proportion of false positives of 26.8%.

The influence of subsetting after TADA analysis on the FDR among selected genes has implications on interpreting the results presented in Figure 1 and Supp. Table 8. At an FDR threshold of 0.2, we identified 34 genes with SPARK damaging variants, including 18 newly significant genes after inclusion of SPARK data. Based on the simulation results, we estimate the expected false positives among 34 genes is about 5 (34*0.159) and among 18 newly emerging genes is about 4 (18*0.27), which closely match the number of genes (4) left after excluding known ASD/NDD genes and newly emerging genes prioritized based on literature evidence.

The results are also relevant in selecting appropriate FDR threshold for declaring significance. Previous studies either used FDR=0.1 as the stringent threshold^7,8^ or used 0.3 as a lenient threshold to implicate more gene in combination of functional data^7^,^105^. In the current study, using a more stringent FDR threshold has limited power in identifying new disease genes, because most known ASD/NDD were identified from previous studies of *de novo* variants, so they tend to have very small q-values in the meta-analysis and are concentrated among most significant genes (Figure 1). Only *BRSK2* and *ITSN1* have FDR<0.1 after the meta-analysis.

We chose not to use a more lenient threshold of 0.3. Although we showed that conditioning on observing variants in SPARK makes the FDR among selected genes slightly conservative, we also noted about half those genes have already been established as ASD/NDD genes. Taking them as true positives, the FDR among new genes will be much higher than 0.3, weakening their statistical evidence. For example, by lowering the FDR threshold to 0.3, we found an additional 9 genes, including one known ASD gene *CACNA1H* (Supp Table 8). Based on the simulation results and accounting for the analysis already done for the 34 genes at FDR threshold of 0.2, we estimated more than half (5.6=43*0.224-4) are likely false positives. Nonetheless, we present some functional evidence supporting *CUL1*, *SRRM2*, and *EPN2* for genes at this FDR threshold in Supp Table 10. We also noted that at the FDR threshold of 0.3, genes affected by only one *de novo* damaging variants will start to pop up (Supp Table 8). Together it suggests that genes identified with an FDR threshold between 0.2 and 0.3 should best be left for further replication in future studies.

Based on the justifications above, an FDR threshold of 0.2 is a reasonable tradeoff to maximize the discovery power in this study without incurring high false positive findings.

## Author contributions

P.F., X.Z., I. Astrovskaya, T.N.T., J.J.M., B.J.O., N.V., Y.S. and W.K.C. **designed and conceived the study**. A.C., A.C.G., A.D.S., A.E., A.G., A.J., A.J.A., A.L.R., A.M., A.M.D., A.N., A.N.S., A.P., A.P.M., A.R.S., A. Swanson, B.A.H., B.E.R., B. Hauf, B.J.O., B.M.V., B.V., C‥A., C.A.E., C.C., C.E.R., C. Harkins, C. Hayes, C.H.W., C.J.S., C. Lord, C.O., C.R.R., C.T., D.E.S., D.G.A., D.I., D. Lee Coury, D. Li, E.A.F., E. Berry-Kravis, E.C., E.J.F., E.L., E.L.W., E.M.B., E.O., E.T.M., G.M., G.S.D., H.E.K., H.H., H. Lam Schneider, H. Lechniak, H. Li, H. Zaydens, I. Arriaga, J.A., J.A.G., J.F.C., J.G., J.L., J.M., J.N., J.O., J.P., J.P., J.S., J.S.S., J.T., J.T.M., J. Wallace, K.A.S., K.C., K.E.H., K.G.P., K.L.P., K.O., K. Roeder, L.A., L.A.C., L. Beeson, L.D., L.D.P., L.G.S., L.M.H., L.M.P., L.P.G., L.S., L.V.S., L. Wasserburg, L. Casey White, L.Y.H., M.A., M.C., M. Heyman, M. Jones, M. Jordy, M.J.M., M.N.H., M.S., M.T., M.Y., N.B., N. Hanna, N. Harris, N. Lillie, N.R., N.T., O.Y.O., P.F., P.M., R.A. Bernier, R.D.A., R.D.C., R.J.L., R.P.G., R. Remington, R.T.S., S.B., S.C., S.E., S.F., S.G., S.H., S.J., S.J.B., S.J.L., S.L.F., S.L.H., S.M.K., S.P., S.Q., S. Sandhu, S.T., S.W., V.J.M., V.S., W.K.C., W.S.Y., and Z.W. **recruited participants and collected clinical data and biospecimens**. A. Balasubramanian, A. Beaudet, A.F., A.H., A.J. Griswold, A.K., A. Soucy, B.J.O., C.L Martin, C.N., D.H.G., E. Berry-Kravis, E. Bahl, E.E.E., E.R., H. Doddapaneni, H.Q., H.Zhang, H.Zhao, I. Astrovskaya, J.B.H., J.H., J.J.M., J.S.S., J.V., J. Wang, K. Rajbhandari, L. Brueggeman, L.G.S., L.P.G., L.Z., M.G., M.R.G., M.Y.D., N.V., P.F., R.A. Barnard, R.D., R.D.A., S.L., S.M., S. Santangelo, S. Skinner, S.X., T.K., T.N.T., T.P., T.S., T.T., T.W., T.Y., V.G., V.K., W.K.C., X.L., X.Z., Y.D., Y.S., and Z.M. **helped with data interpretation**. A.A., A. Bashar, A.E.L., A. Salomatov, A.S.C., B. Han, C.M.S., C.R., I. Astrovskaya, I.F., J.A., J.B.H., K.L., M.D.M., M.E.B., M.K., N.C., N. Lawson, N.V., R.M., R. Rana, S.G., S. Shah, S.X., W.C., and W.J. **built and supported the SPARKforAutism.org website, software, databases and systems, and managed SPARK data**. A.D.K., A.H., A.J. Gruber, A. Sabo, B.J.O., C.E., D.M., E. Brooks, G.J.F., I. Astrovskaya, J.J.M., L. Brueggeman, L.G.S. M.P., M.R.G., N.V., P.F., R.A. Barnard, R.A.G., T.N.T., T.W., W.K.C., X.Z., and Y.S. **performed analyses, processed biospecimens and sequenced DNA samples.**P.F. and W.K.C. **supervised the work.**P.F., X.Z., I. Astrovskaya, T.N.T., J.J.M., B.J.O., N.V., W.K.C., and Y.S. **wrote this paper.**

## Acknowledgements

We are extremely grateful to the thousands of individuals and families who are participating in this study. We are grateful to the many autism advocacy and service organizations that have helped us inform the community about SPARK, including the Autism Society of America and its affiliates, Autism Speaks, Autism Science Foundation, Easter Seals, Arkansas Autism Resource and Outreach Center, Global and Regional Asperger’s Syndrome Partnership, Kentucky Autism Training Center, and Autistic Self Advocacy Network. We thank the members of SPARK’s Community Advisory Council for providing feedback and advice. We thank members of our Scientific and Community Advisory Board and SFARI scientists for advice on our protocol and participant outreach and retention strategies. We thank PreventionGenetics for managing and processing biospecimens, DNA Genotek for handling saliva kit logistics, and Baylor College of Medicine Human Genome Sequencing Center for exome sequencing. The SPARK initiative is funded by the Simons Foundation as part of SFARI.

This research was supported, in part, by a grant from the National Institute of Mental Health (NIMH R01MH101221) and a grant from the Simons Foundation (SFARI #608045) to E.E.E., a grant from the National Institute of Mental Health to T.N.T. (1K99MH117165) and grants MH105527 and DC014489 from the National Institute of Health to L.B. and J.J.M.

E.E.E. is an investigator of the Howard Hughes Medical Institute. B.J.O. is a Klingenstein-Simons Fellow (Esther A. & Joseph Klingenstein Fund, Simons Foundation).

B.J.O. is a Klingenstein-Simons Fellow (Esther A. & Joseph Klingenstein Fund, Simons Foundation)

