## Supplementary material for "Exome sequencing of 457 autism families recruited online provides evidence for novel ASD genes": Supp Figures and Supp Table 8

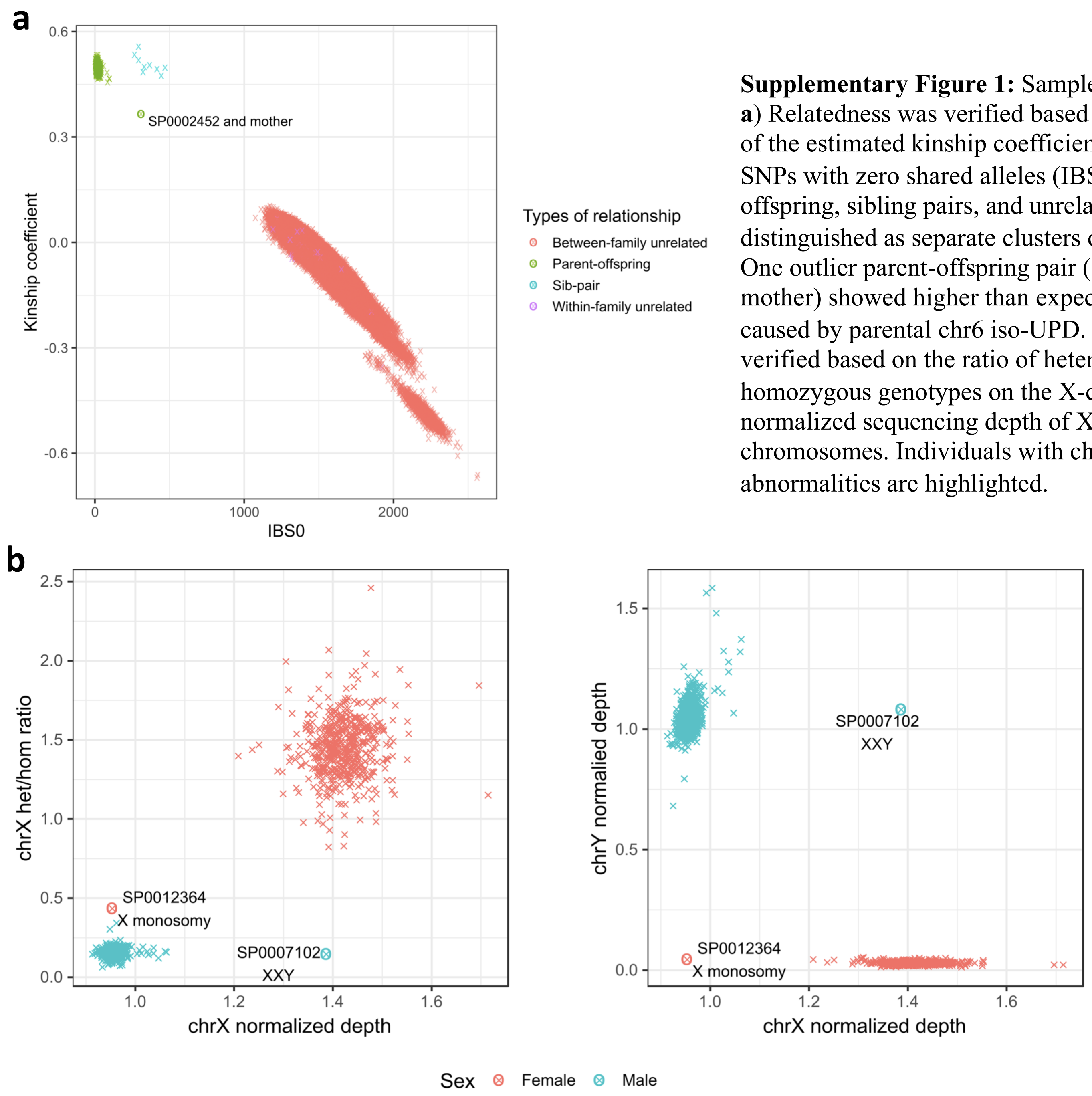


**Supplementary Figure 1:** Sample quality controls. **a**) Relatedness was verified based on the scatterplot of the estimated kinship coefficient and number of SNPs with zero shared alleles (IBS0). Parent-offspring, sibling pairs, and unrelated pairs can be distinguished as separate clusters on the scatterplot. One outlier parent-offspring pair (SP0002452 and mother) showed higher than expected IBS0 and was caused by parental chr6 iso-UPD. **b**) Sample sex was verified based on the ratio of heterozygous to homozygous genotypes on the X-chromosome, using normalized sequencing depth of X and Y chromosomes. Individuals with chromosomal abnormalities are highlighted.

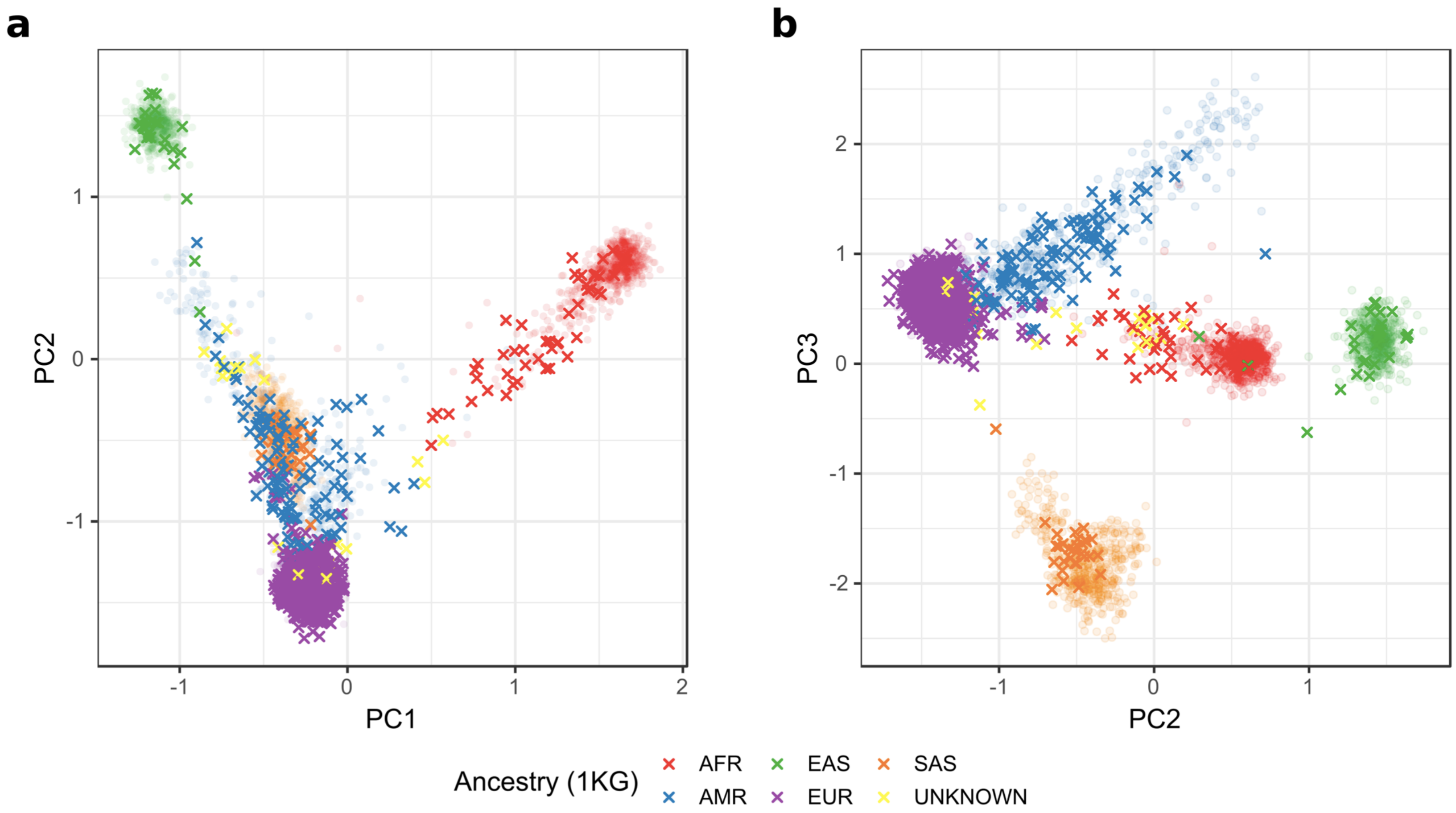


**Supplementary Figure 2:** Principal component (PC) analysis of sample ethnicity. Samples were projected onto the PC axes defined by the samples from 1000 Genomes Project (shown in light colors). **a**) The first two PCs can distinguish samples from three major continents. **b**) PC3 further distinguishes South Asians from Admixed Americans. Sample ethnicities were inferred based on the first four PCs using a machine learning approach implemented in peddy [Pedersen AJHG 2017].

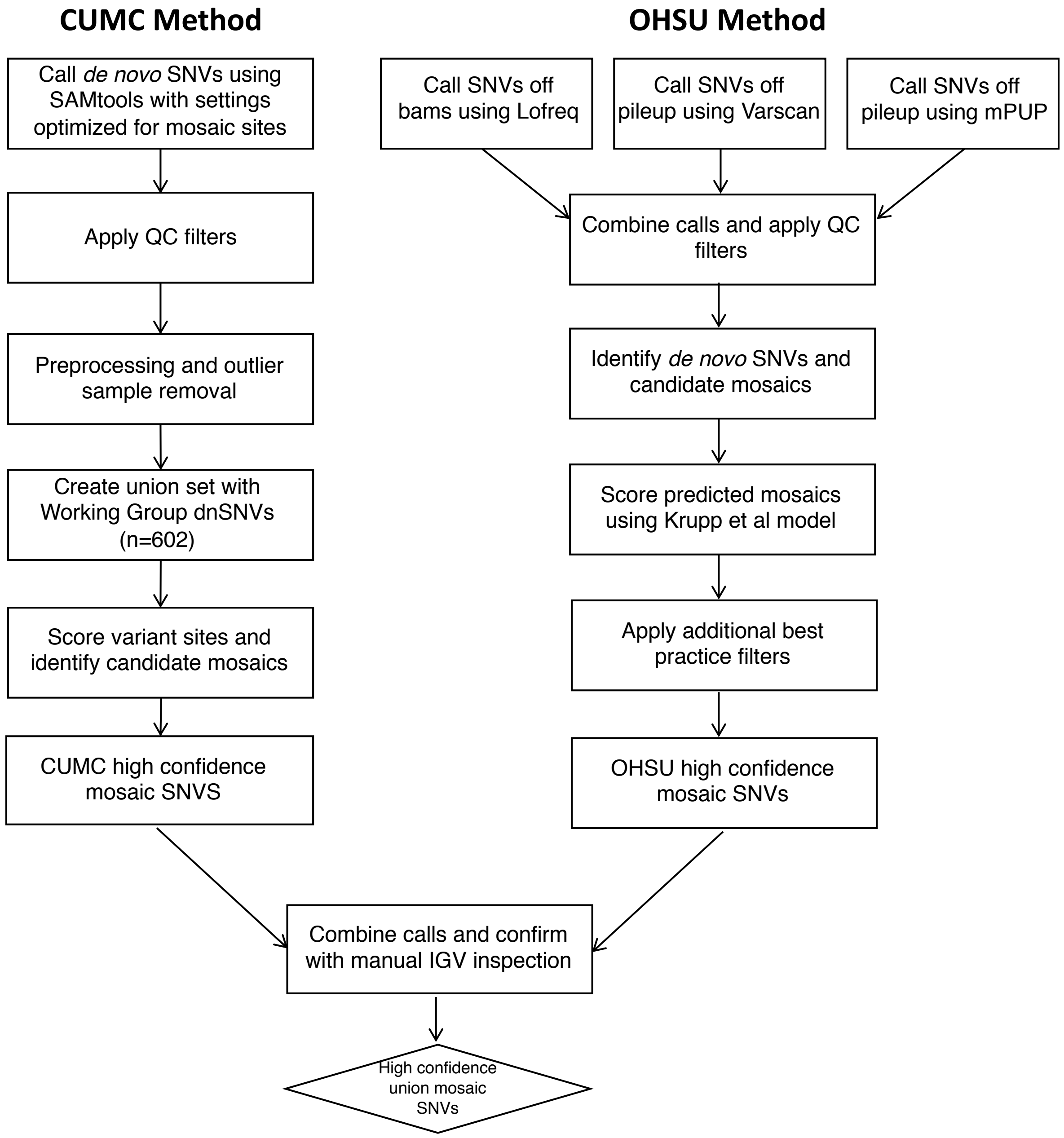


**Supplementary Figure 3:** Parallel calling approach for mosaic SNVs.

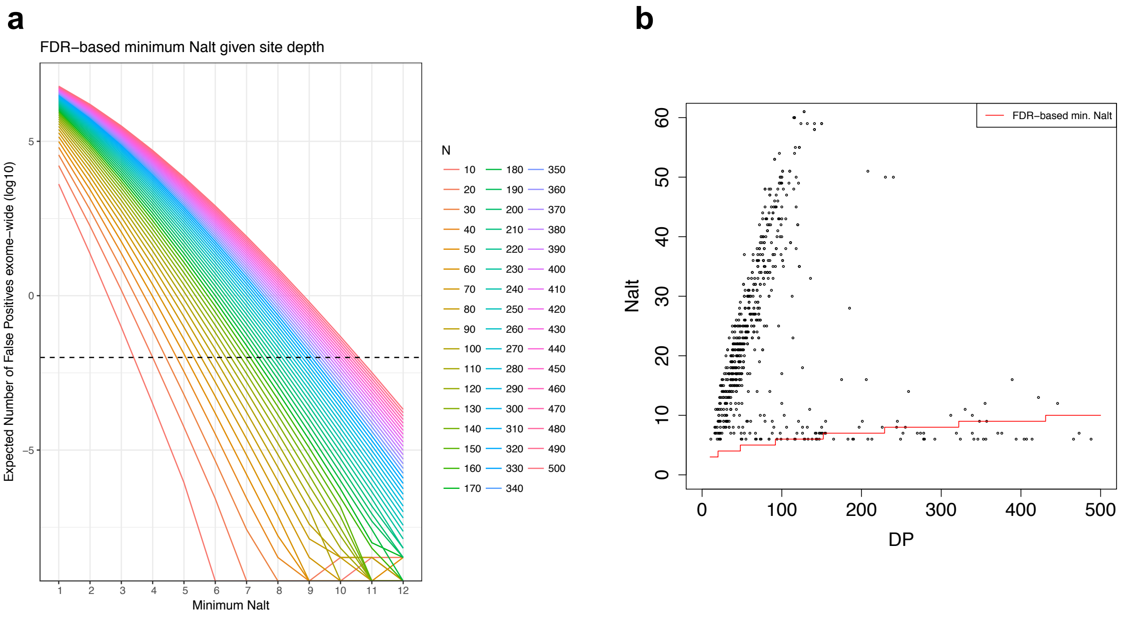


**Supplementary Figure 4:** CUMC Method development: FDR-based minimum Nalt threshold. Data shown are variants called by samtools. **a)** Theoretical FDR-based minimum alternate allele read depth (Nalt) thresholds as a function of total read depth (N). Assuming that sequencing errors are independent and that errors occur with probability 0.005, with the probability of an allele-specific error being 0.005/3=0.00167, and given the total number of reads (N) supporting a variant site, we iterated over a range of possible Nalt values between 1 and 0.5*N and estimated the expected number of false positives due to sequencing error, exome-wide [(1-Poisson(Nalt, $\lambda$=N*(0.00167))* $3\times10^7$ ]. Assuming one coding de novo SNV per individual[58] and that roughly 10% of de novo SNVs arise post-zygotically[20-22], we estimate there to be 0.1 mosaic mutations per exome. Under this assumption, to constrain theoretical FDR (in terms of distinguishing low allele fraction sites from technical artifacts) to 10%, we allowed a maximum of 0.01 false positives per exome. We used this cutoff to identify a FDR-based minimum Nalt threshold for each site as a function of total site depth. The dashed line denotes the threshold at which the expected number of false positives exome-wide is 0.01. **b)** FDR-based minimum Nalt threshold applied to samtools calls. Variant calls are plotted using total read depth (DP) and alternate allele read depth (Nalt). The red line marks the Nalt cutoff as a function of DP.

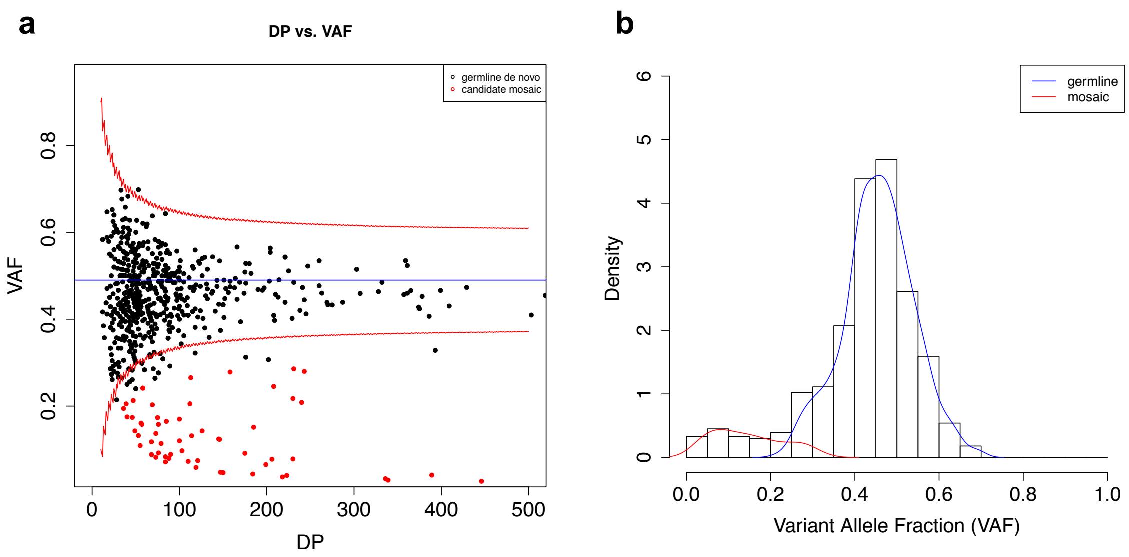


**Supplementary Figure 5:** CUMC Method development: mosaic candidate identification. Data shown are the consensus variant calls. **a)** Total read depth (DP) in relation to variant allele fraction (VAF). The blue line denotes the Beta-Binomial mean VAF and the red lines denote the 95% confidence interval. To calculate the posterior odds that a given variant arose post-zygotically, we first calculated a likelihood ratio (LR) using two models: M0: germline heterozygous variant, and M1: mosaic variant. Under our null model M0, we calculated the probability of observing Nalt from a beta-binomial distribution with site depth N, observed mean germline VAF p, and overdispersion parameter θ. Under our alternate model M1, we calculated the probability of observing Nalt from a beta-binomial distribution with site depth N, observed site VAF p=Nalt/N, and overdispersion parameter θ. Finally, for each variant, we calculated LR by using the ratio of probabilities under each model and posterior odds by multiplying LR by our E-M estimated prior mosaic fraction estimate. Sites with posterior odds greater than 10 were predicted mosaic (corresponding to 9.1% FDR).
**b)** Expectation-Maximization (EM) decomposition of variant allele fraction (VAF) into germline and mosaic distributions. Blue and red lines denote smoothed density curves for each distribution. We used an expectation-maximization (E-M) algorithm to jointly estimate the fraction of mosaics among apparent de novo mutations and the false discovery rate of candidate mosaics. This initial mosaic fraction estimate gives a prior probability of mosaicism independent of sequencing depth or variant caller and allows us to calculate, for each variant in our input set, the posterior odds that a given site is mosaic rather than germline.

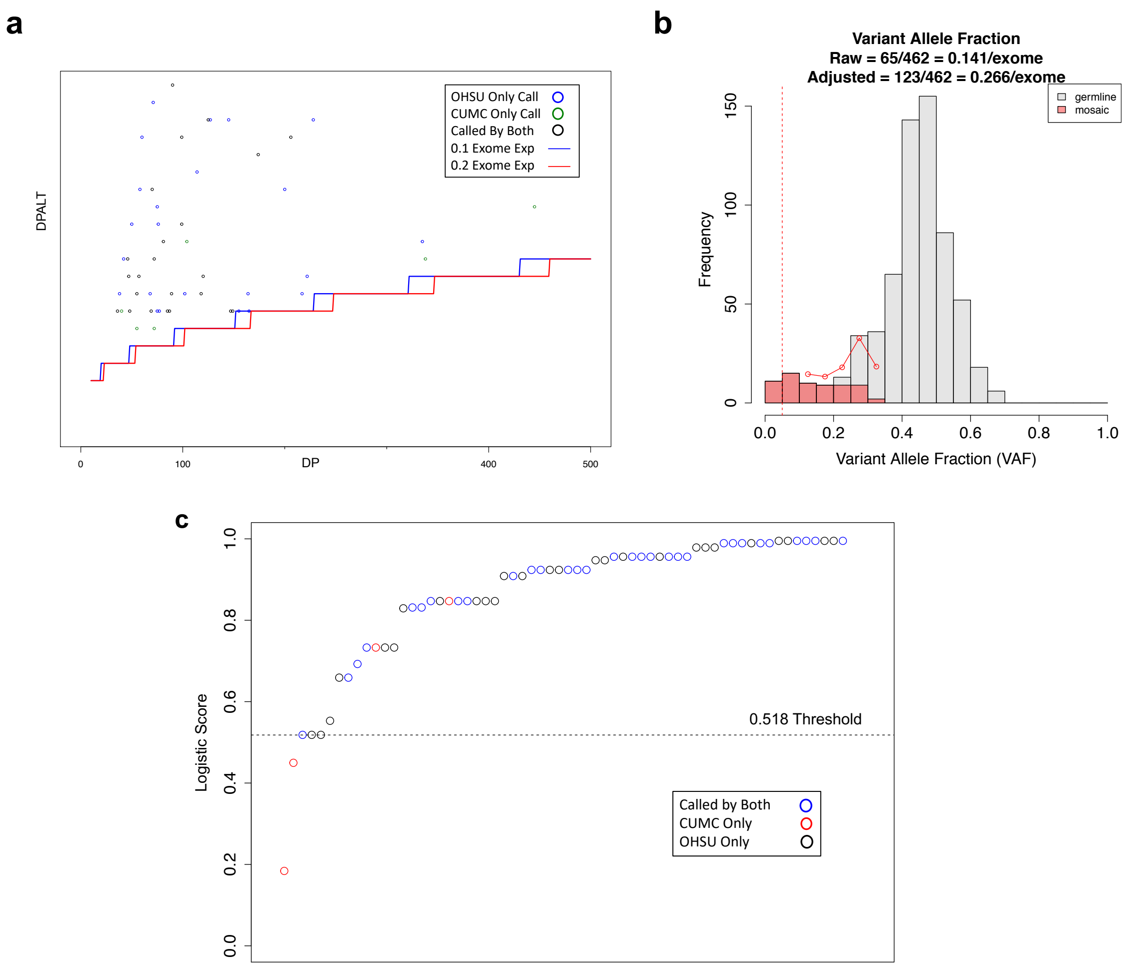


**Supplementary Figure 6:** Characterization of high confidence union mosaic calls. **a)** Alternative allele depth in relation to FDR based threshold. All calls are above the FDR threshold for both a 0.1 or 0.2 events per exome expectation. **b)** Variant allele fraction distribution. The grey and red bars denote germline and mosaic variants, respectively. The red line denotes the estimated true number of mosaics at each VAF window adjusted for mosaic detection power. Detection power is estimated as a function of variant allele fraction and sample average sequencing depth. The dashed vertical line denotes 5% VAF, below which estimated detection power is extremely limited and likely to artificially inflate adjusted counts. **(c)** Percentile ranked distribution of Krupp et al.[22] logistic mosaic score, 0.518 was the applied threshold for OHSU pipeline. Scores are overall well distributed between overlapping and group specific calls. Three of the CUMC only calls were not scored as they were filtered out of the OHSU pipeline before scoring due to differences in segmental duplication annotation.

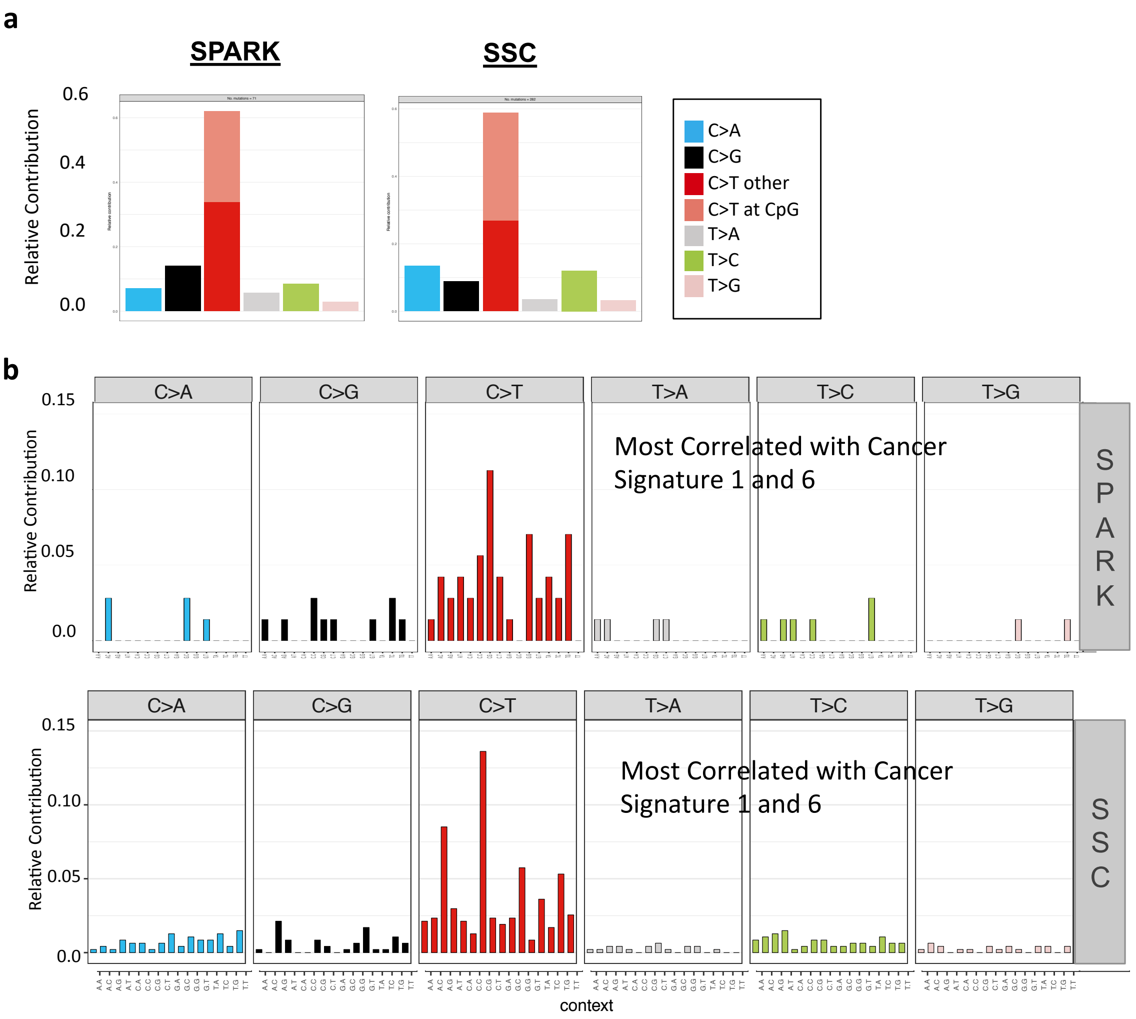


**Supplementary Figure 7:** Comparison of Mosaic Mutations Spectra and Signatures in SPARK and SSC. Mutational contexts and frequency were extracted and plotted using the R package MuationalPatterns[59]. **a)** Mutational spectrum of the six different possible substitutions for SSC and SPARK mosaic mutations. **b)** Mutational signature of the relative frequency of mutations (Y-axis) within trinucleotides (context) for SSC and SPARK mosaic mutations. Though there are fewer calls in SPARK due to the smaller cohort size, both SSC and SPARK show a strong correlation to the same Cancer Signatures which are indicative of endogenous and DNA mismatch repair mutational processes.

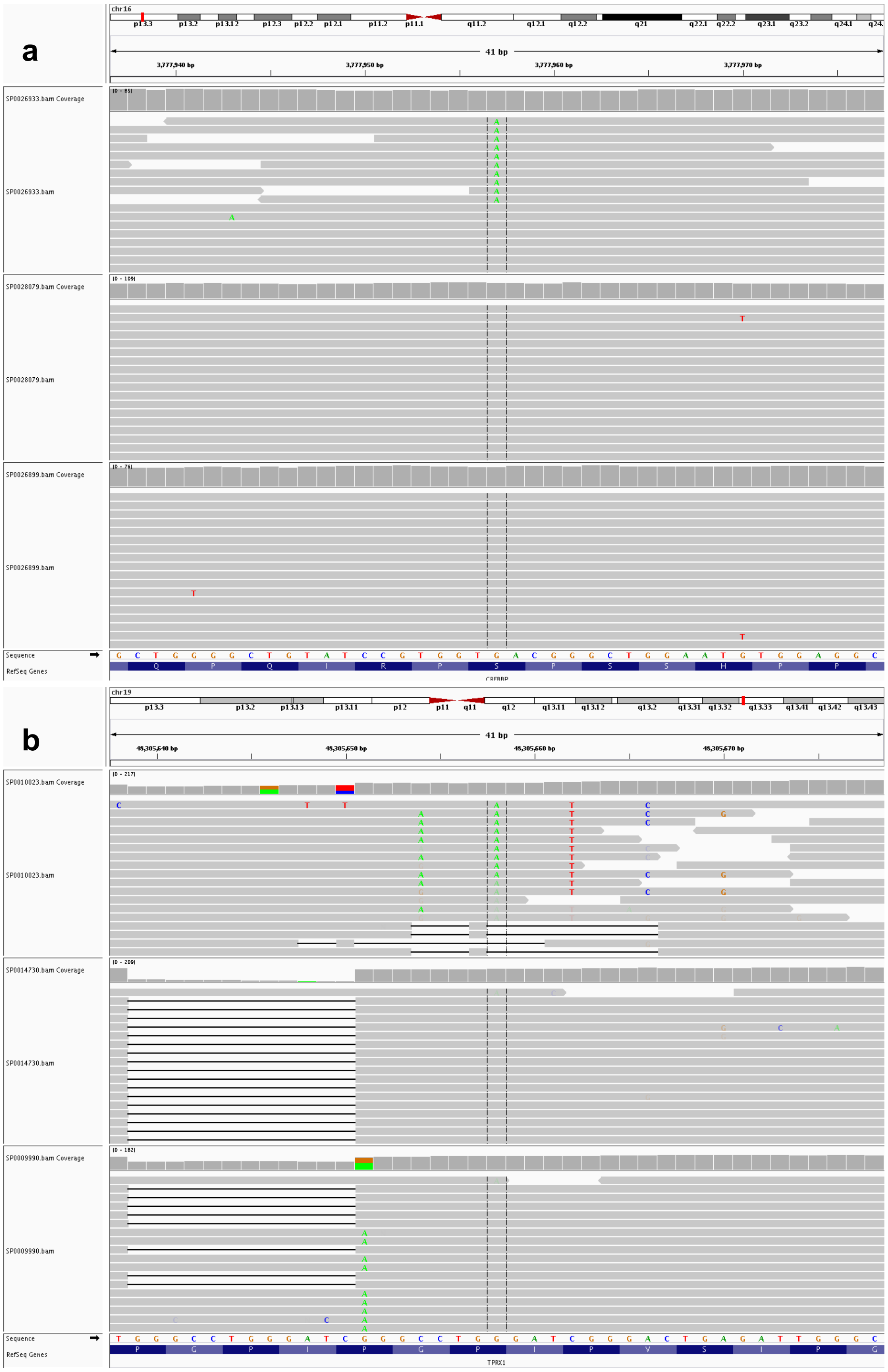


**Supplementary Figure 8:** IGV plots used in mosaic mutation visualization and review. **a)** Example mosaic candidate passing IGV review – SP0026933:chr16:3777957:G>A. **b)** Example mosaic candidate failing IGV review – SP0010023:chr19:48305658:G>A.

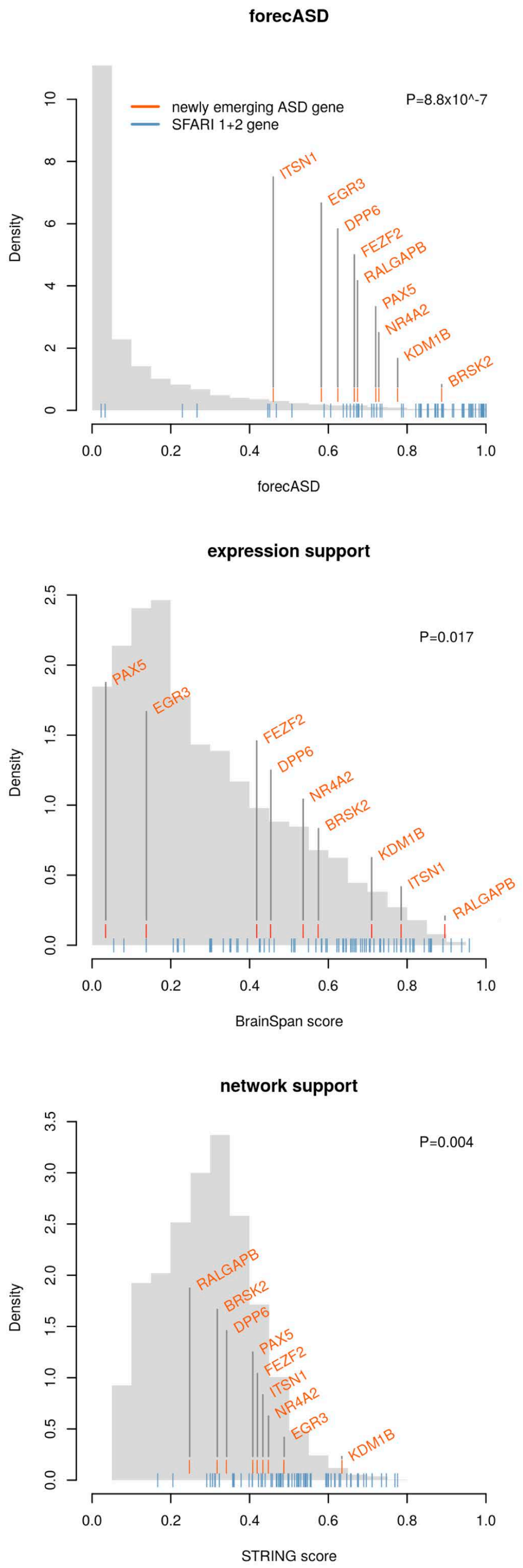


**Supplementary Figure 9:** Support for newly emerging ASD genes. The emerging genes highlighted in this work have significantly elevated forecASD scores (P=8.8x10-7 ,Wilcoxon test), with all genes in the top decile. Two constituent features in the forecASD ensemble (brain spatiotemporal expression and network topology) also show significantly elevated scores (P=0.017 and P=0.004, respectively), suggesting that these genes show similar properties to known ASD genes beyond genetic association and across a diverse feature space, thereby supporting the robust biological plausibility of these genes.

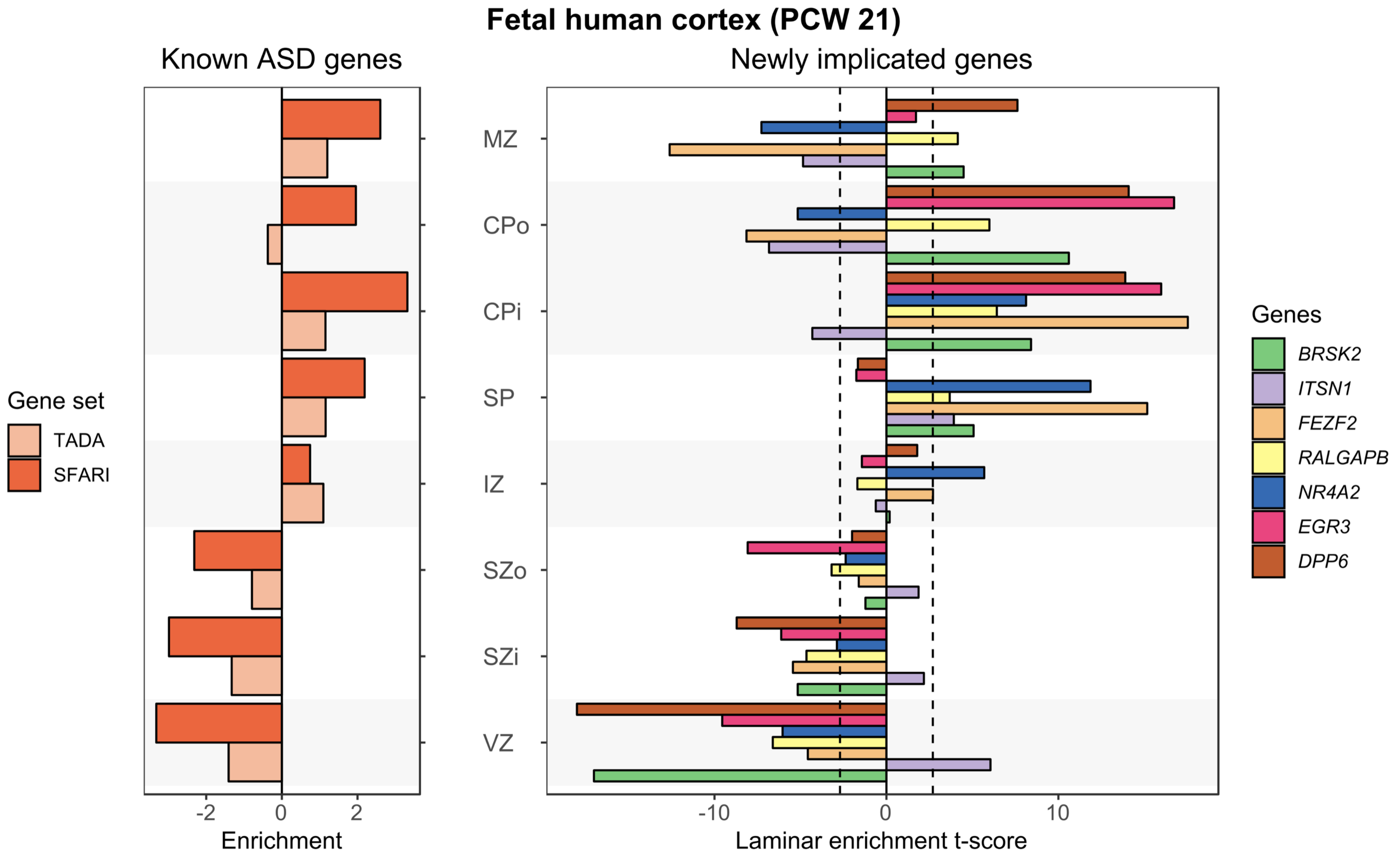


**Supplementary Figure 10:** Gene expression of newly emerging ASD genes in human fetal brain PCW21.

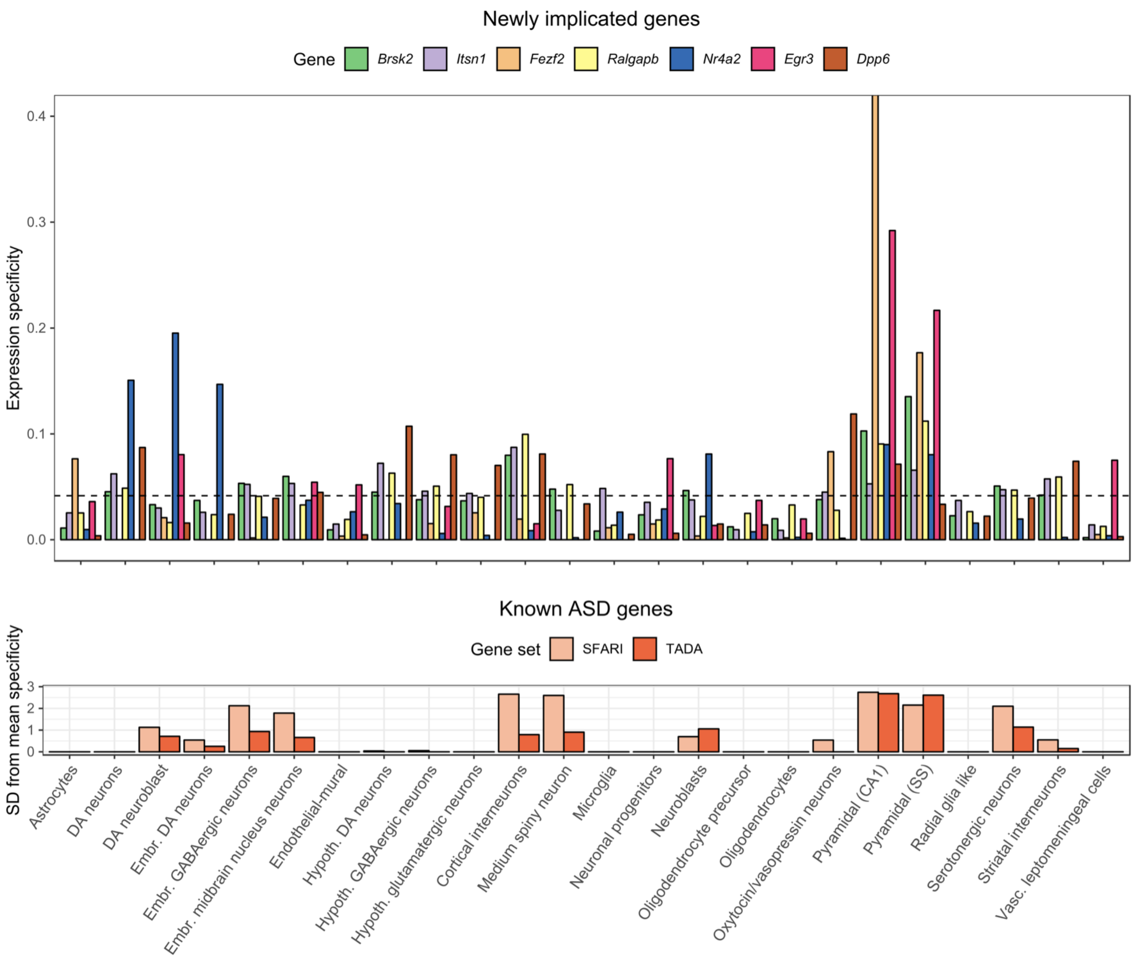


**Supplementary Figure 11**: Expression specificity of newly emerging ASD genes in single-cell RNA-seq data from fetal and adult mouse brains. The specificity of expression in a cell type is measured by a specificity index which is the mean expression level in one cell type over the summation of mean expression level across all cell types[61]. For a gene set, the mean expression specificity of its genes was compared with 10,000 sets of randomly drawn genes matched for the transcript length and GC content and the enrichment is measured by the standard deviation from the mean specificity of random gene sets[61]. The mouse neuronal cell types are defined by the analysis of single cell RNA-seq data of fetal and adult mouse brains generated by Karolinska Institutet (KI) and used in a previous study[48]. The mouse orthologs of human genes were retrieved from MGI database[62]. The known ASD genes show highest enricment in pyramidal neurons (in hippocampus CA1 and somatosensory cortex), cortical interneurons, and medium spiny neurons. The first three enriched cell types were previously reported for the 65 autism genes identified from TADA meta-analysis[61]. The newly implicated genes also show highest specificity in pyramidal neurons, suggesting functional convergence in these cell types.

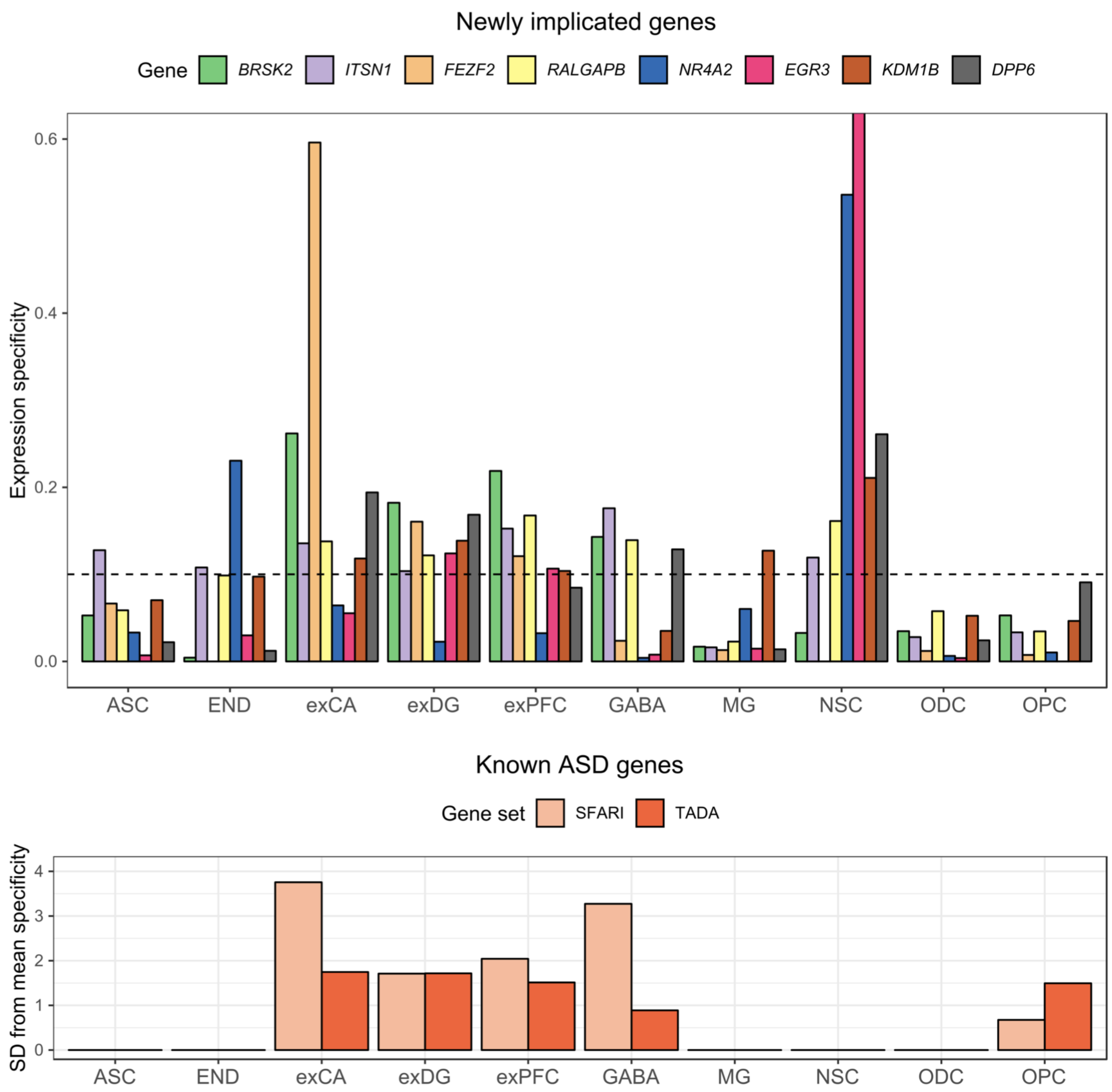


**Supplementary Figure 12:** Expression specificity of newly emerging ASD genes in single-cell RNA-seq data from human brains. Human neuronal cell types are defined by the single-nucleus RNA-seq data of archived human brains[49]. Known and new ASD genes were mostly enriched in neurons (exCA, exDG, exPFC) and interneurons (GABA). Highest enrichment was also observed in pyramidal neurons (exXCA). New ASD genes were also enriched in neuronal stem cells that are not implicated by known ASD genes, but the enrichment is not significant. Significance code: *= $p<0.01$, **= $p<0.001$. exPFC=glutamatergic neurons from the PFC, exCA=pyramidal neurons from the hippocampus CA region, GABA=GABAergic interneurons, exDG=granule neurons from the Hip dentate gyrus region, ASC=astrocytes, NSC=neuronal stem cells, MG=microglia, ODC=oligodendrocytes, OPC=oligodendrocyte precursor cells, NSC=neuronal stem cells, SMC=smooth muscle cells, END=endothelial cells.

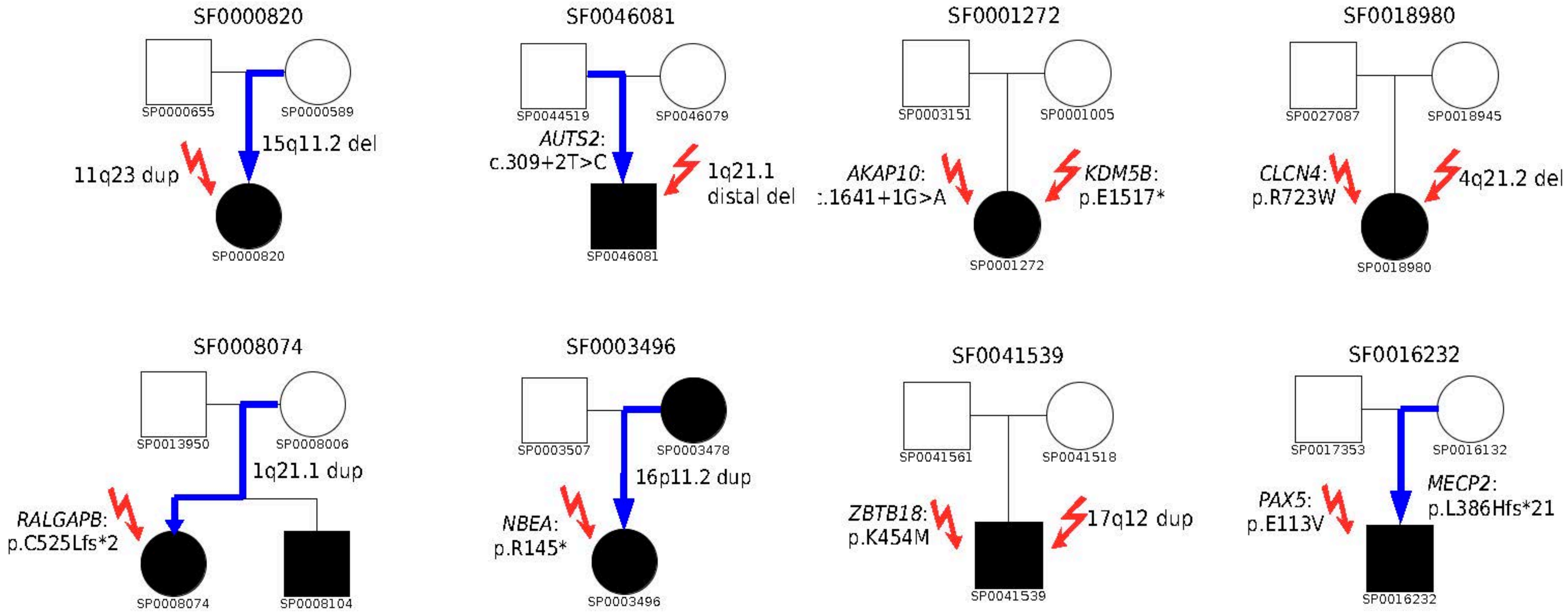


**Supplementary Figure 13:** Eight pedigrees with multiple contributing variants.

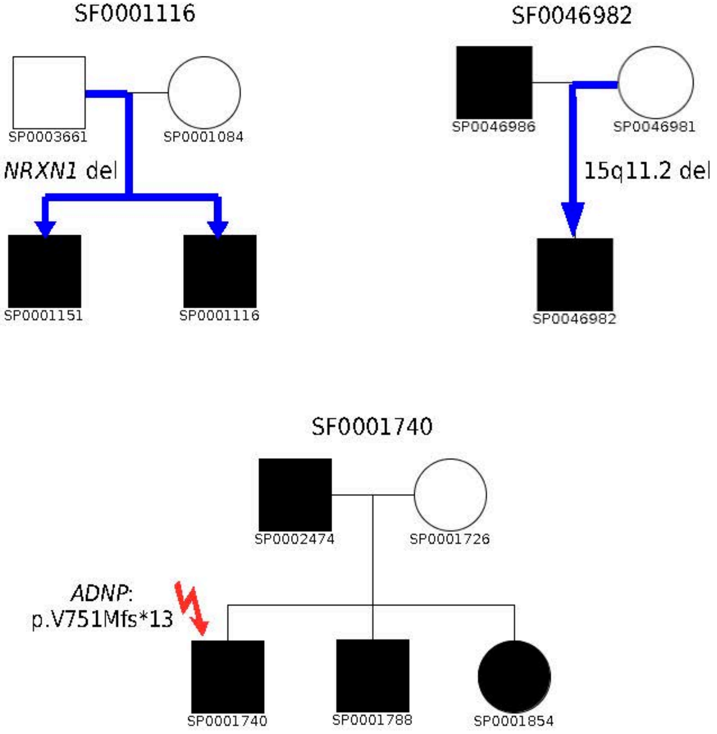


**Supplementary Figure 14:** Genetic causes of ASD were identified in 6 offspring in 5 multiplex families, 3 of which are shown in this figure. SF0003496 and SF0008074 are shown in Supplementary Figure 13. SF0003496 has another affected offspring, who was not sequenced in this study.

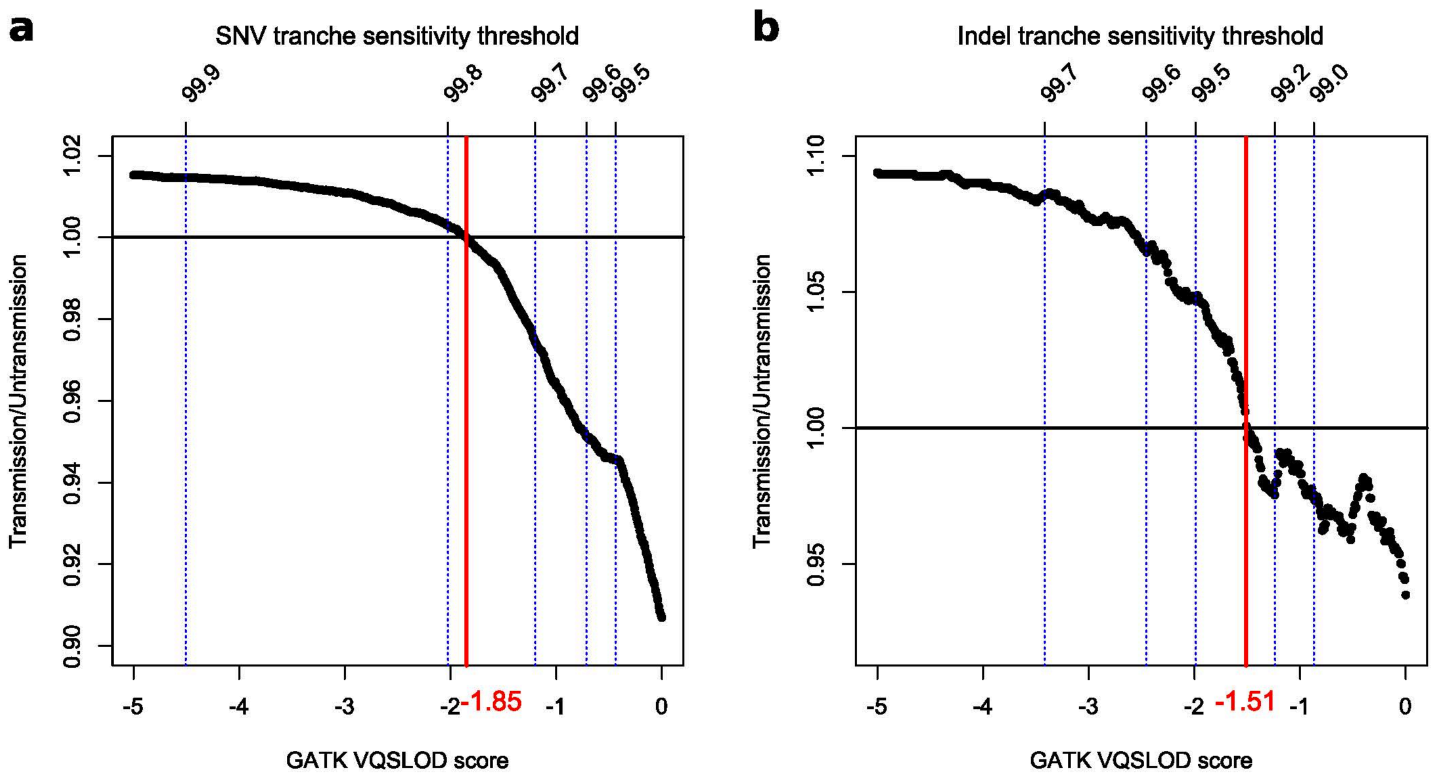


**Supplementary Figure 15:** Recalibrating VQS LOD threshold for analyzing inherited singleton variants. The transmission to un-transmission ratio of singleton synonymous SNVs (**a**) and non-frameshift indels (**b**) are shown as a function of the VQS LOD score. The dashed lines mark the GATK defined cutoffs based on different tranche sensitivity thresholds. The red line shows the cutoffs that balance the transmission to un-transmission ratio and were used in filtering singleton variants for transmission disequilibrium analysis.

| Gene selection method | 1. q-value<=FDR threshold and have contributing de novo variants in a subset of 465 trios | | | 2. q-value>FDR threshold in 4773 trios, and <=FDR threshold after inclusion of 465 additional trios | | |
|---|---|---|---|---|---|---|
| FDR threshold | Total positives | True positives | %FP | Total positives | True positives | %FP |
| 0.1 | 33.50 | 31.05 | 7.3% | 18.38 | 16.10 | 12.4% |
| 0.2 | 45.04 | 37.87 | 15.9% | 25.05 | 18.35 | 26.8% |
| 0.3 | 53.39 | 41.42 | 22.4% | 31.4 | 20.08 | 36.1% |

**Supplementary Table 8:** TADA analysis on simulated data of 5,238 trios. The number of positive findings, true positives, and fraction of false positives (FR) at different FDR thresholds are averaged over 200 repeats.
